# Engineered small extracellular vesicles as a FGL1/PD-L1 dual-targeting delivery system for alleviating immune rejection

**DOI:** 10.1101/2021.06.14.448282

**Authors:** Hsiang-i Tsai, Yingyi Wu, Xiaoyan Liu, Zhanxue Xu, Longshan Liu, Changxi Wang, Huanxi Zhang, Yisheng Huang, Linglu Wang, Weixian Zhang, Dandan Su, Fahim Ullah Khan, Xiaofeng Zhu, Rongya Yang, Yuxin Pang, John E. Eriksson, Haitao Zhu, Dongqing Wang, Bo Jia, Fang Cheng, Hongbo Chen

## Abstract

There is an urgent need for developing new immunosuppressive agents due to the toxicity of long-term use of broad immunosuppressive agents post organ transplantation. Comprehensive sample analysis revealed dysregulation of FGL1/LAG-3 and PD-L1/PD-1 immune checkpoints in allogeneic heart transplantation mice and clinical kidney transplant patients. In order to enhance these two immunosuppressive signal axes, we developed a bioengineering strategy to simultaneously display FGL1/PD-L1 (FP) on the surface of small extracellular vesicles (sEVs). Among various cell sources, FP sEVs derived from mesenchymal stem cells (MSCs) not only enriched FGL1/PD-L1 expression but also maintained the immunomodulatory properties of unmodified MSC sEVs. Next, we confirmed that FGL1 and PD-L1 on sEVs were specifically bound to their receptors LAG-3 and PD-1 on target cells. Importantly, FP sEVs significantly inhibited T cell activation and proliferation *in vitro* and a heart allograft model. Furthermore, FP sEVs encapsulated with low-dose FK506 (FP sEVs@FK506) exerted stronger effects on inhibiting T cell proliferation, reducing CD8^+^ T cell density and cytokine production in the spleens and heart grafts, inducing regulatory T cells in lymph nodes, and extending graft survival. Taken together, dual-targeting sEVs have the potential to boost the immune inhibitory signalings in synergy and slow down transplant rejection.

## 1. Introduction

Immunosuppressants such as calcineurin inhibitor FK506 (Tacrolimus) or cyclosporin A (CsA) are widely used in organ transplantation^[1]^. Their prolonged use, however, can result in a multitude of side effects, including nephrotoxicity, infection, bone marrow suppression and gastrointestinal reactions^[2]^. There is therefore an unmet need for the development of drugs based on novel mechanisms and targets to replace, or use in concert with, conventional immunosuppressive drugs, to enhance the overall survival rate of patients.

T-cell expressed immune checkpoints (ICPs) such as PD-1, TIM-3, VISTA, TIGIT, and LAG-3 interact with their inhibitory ligands expressed on antigen-presenting cells and/or on tumor cells, responsible for maintaining self-tolerance and preventing an autoimmune response^[3]^. Immune checkpoint inhibiting therapeutic antibodies have been recently reported as an effective strategy for blocking these immunosuppressive axes, thereby overcoming tumor immune evasion^[4]^. Bispecific antibodies are also important in this regard and have been demonstrated to interrupt a number of these axes (e.g. CTLA-4/CD80, PD-1/PD-L1), resulting in excellent therapeutic effects against malignant tumors^[5]^. Given the important role of ICPs in immune tolerance, it is reasonable to use ICPs as a group of therapeutic targets to restore immune tolerance and treat organ transplant rejection or autoimmune diseases. However, it is technically very challenging to develop ICP agonist antibodies, especially bispecific ICP agonist antibodies to restore immune tolerance^[6]^.

In recent years, cell membrane-based nanovesicle delivery systems that readily display transmembrane proteins such as ICPs, have seen substantial development^[7]^. We previously engineered cell membrane-derived nanovesicles (NVs) displaying PD-L1/CTLA-4 dual-targeting cargos. These NVs consequently enhanced PD-L1/PD-1 and CTLA-4/CD80 immune inhibitory pathways, exerted immune inhibitory effects, and prolonged the survival of mouse skin and heart grafts^[8]^. However, CTLA-4 binds to CD80 or CD86 on antigen-presenting cells (APCs) rather than T cells and operates at different stages and locations of immune inhibition than the PD-L1 pathway^[9]^. Furthermore, NVs are generated by serial extrusion of the cell membrane, leading to the membrane incompleteness, inner and outer membrane turnover, and incorrect arrangement of membrane molecules during preparation^[8]^. In contrast, small extracellular vesicles (sEVs), a 50–150 nm membrane vesicle containing miRNAs and proteins, have been identified as a superior alternative to natural membrane delivery systems^[10]^ largely due to their high biocompatibility and negligible side effects. Interestingly, melanoma cells are reported to secrete sEVs carrying a high level of PD-L1 and effectively suppress CD8^+^ T cell activity^[11]^. Thus, combinational inhibition of both PD-1 and other ICPs on T cells by using sEVs simultaneously carrying multiple target ligands might be a more potent therapeutic strategy for attenuating T-cell mediated immune rejections.

Mesenchymal stem cells (MSCs) have been increasingly used for autoimmune disease treatment due to their immune modulation functions^[12]^. Recently, it was determined that MSCs exert their immune-modulatory effects, mostly by secreting soluble factors and sEVs^[13]^. MSC derived sEVs have been reported to have therapeutic effects after organ transplantation and Graft-versus-host disease (GvHD), including immunosuppression, anti-inflammatory properties, and the induction of tissue regeneration^[14]^. These favorable characteristics underscore the potential of MSC-sEVs as promising target vehicles to foster immune tolerance.

In this study, we first identified a simultaneous rise of PD-1 and LAG-3, but not their partner proteins PD-L1 and FGL1 in heart transplantation models and clinical kidney transplant patients. We then bioengineered MSCs to obtain sEVs simultaneously displaying highly surface FGL1/PD-L1, and confirmed their specificity to bind their ligands LAG-3 and PD-1 on T cells. The FGL1/PD-L1 dual-targeting sEVs (FP sEVs) inhibited the activation of T cells both *in vitro* and in a mouse heart transplantation model, via enhancing the immunosuppressive pathways of both FGL1/LAG-3 and PD-L1/PD-1. Importantly, FP sEVs encapsulated with FK506 displayed stronger inhibition of T cell proliferation than FP sEVs or FK506 alone *in vitro* and *in vivo*, providing strong evidence for a synergistic effect between low-dose FK506 and sEVs expressing FGL1/PD-L1 leading to the weakened alloimmune response and induced allograft tolerance. This constitutes a novel strategy for the development of multi-targeting genetically engineered sEVs as effective immunosuppressants to inhibit post-transplant rejection.

## 2. Results

### 2.1 LAG-3/PD-1 and FGL-1/PD-L1 expression levels are inconsistent in organ transplant recipients experiencing rejection

In order to demonstrate if activating inhibitory axes could be advantageous for the relief of immunologic rejection, we first checked mRNA expression levels among a panel of immune checkpoint genes in a heterotopic heart allograft mouse model. We found that among these checkpoint genes, *Lag-3* and *Pd-1* mRNA in the spleens were simultaneously upregulated 8.56- and 6.24-fold higher in the transplantation group (**Figure 1A**). We then further checked *LAG-3* and *PD-1* expression in peripheral blood mononuclear cells (PBMCs) of clinical renal transplant patients. We found that *LAG-3* and *PD-1* expression levels in patients with T-cell mediated rejection (TCMR) were significantly higher than in both antibody-mediated rejection (ABMR) recipients and non-rejection recipients after kidney transplantation (**Figures 1B-C**). Next, to identify the expression of *Lag-3* and *Pd-1* in T cell subsets for organ transplant rejection, the CD8^+^ T cells and CD4^+^ T cells were isolated from the spleens of mouse heart transplantation model by kits, then the expression levels of *Lag-3* and *Pd-1* were detected by qPCR. As showed in **Figures 1D-E**, it was found that with the occurrence of transplantation rejection, the expression levels of *Lag-3* and *Pd-1* increased obviously in CD4^+^ T cells, but more significantly in CD8^+^ T cells. All of the above results indicate that when immune rejection occurs, T cells attempt to brake the over-activated CD8^+^ T cells and re-establish immune tolerance by upregulation of PD-1 and LAG3. Surprisingly, neither PD-1 ligand PD-L1 nor the newly discovered inhibitory ligand of LAG-3, FGL1 increased in the spleens of transplanted mice (**Figure 1F**). Thus, we speculated that the simultaneous enhancement of FGL1 and PD-L1 inhibitory molecules would be a potential strategy to activate the PD-L1/PD-1 and FGL1/LAG-3 inhibitory axes, thereby inhibiting CD8^+^ T cell activation and re-establishing immune tolerance of the graft.

**Figure 1.**
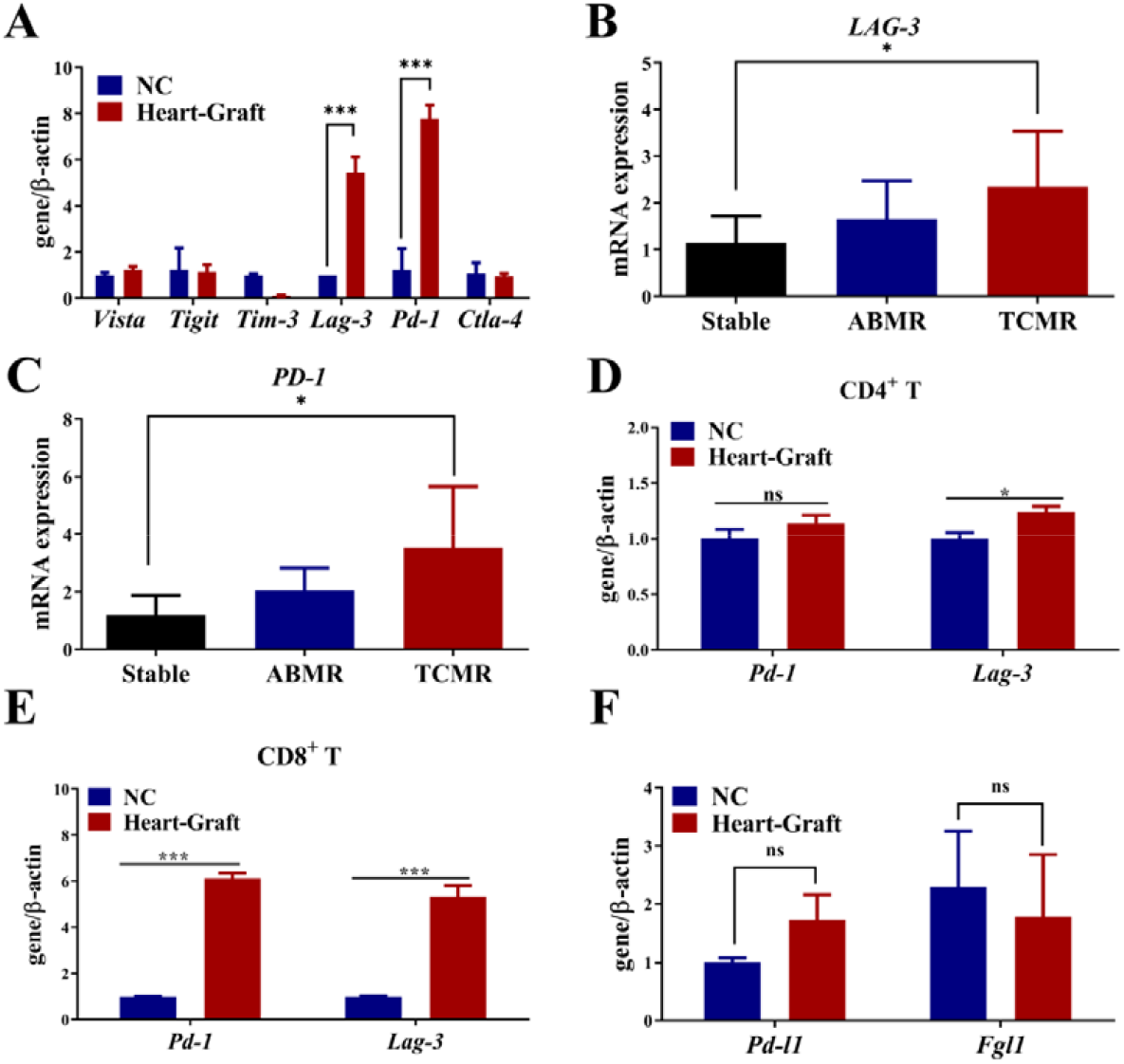
LAG-3/PD-1 is inconsistent with FGL1/PD-L1 in organ transplant recipients with rejection. A) Quantitative PCR to verify the expression of immune checkpoint receptors in the spleens of heart-graft models. NC group: normal mice without heart grafts, *n* = 3. Error bar, mean ± SEM. *P*-values are calculated using student T-test. B-C) Quantitative PCR to verify the expression of *LAG-3* (B) and *PD-1* (C) in PBMCs of clinical renal transplant patients, *n* = 6-7. Error bar, mean ± SEM. *P*-values are calculated using one-way ANOVA by Tukey post-hoc test. D-E) Quantitative PCR to verify the expression of *Pd-1* and *Lag-3* on CD4^+^ T (D) and CD8^+^ T (E) cells were harvested by isolation kit from the spleens of normal and heart-graft mice. NC group: without heart grafts in a heart graft model, *n* = 3. Error bar, mean ± SEM. *P*-values are calculated using student T-test. F) Quantitative PCR verified the ligand expressions of *Fgl1* and *Pd-l1* in the spleens of mouse heart transplant models, NC group: normal mice without heart grafts, *n* = 5. Error bar, mean ± SEM. *P*-values are calculated using student T-test. NS: no significant, \**P* < 0.05, \*\**P* < 0.01, \*\*\**P* < 0.001.

### 2.2 Establishment and Characterization of FGL1/PD-L1 Dual-Targeting sEVs

Recently, small extracellular vesicles (sEVs) released by metastatic melanomas carrying PD-L1 on their surface were reported to successfully suppress the function of CD8^+^ T cells^[15]^, which provided us with a rationale for developing FGL1 and PD-L1 dual-targeting sEVs as an anti-rejection therapy. As the expression of FGL1 and PD-L1 in HEK-293T, HepG2, A549, and MSC cell lines were inconsistent (**Figure 2A**), simultaneous high expression of both FGL1 and PD-L1 on sEVs from the same natural cell line would be difficult. Furthermore, FGL1 and PD-L1 expression in purified sE<u>V</u>s were low and fluctuated among four different cell lines (**Figure 2B**). Thus we speculated that bioengineering sE<u>V</u>s that co-expressed both FGL1 and PD-L1 to a relatively high degree may serve as a strategy to relieve T-cell mediated graft rejection. As FGL1 is a soluble protein and may not be expressed on the membrane of sEVs, we first reconstructed a membrane-localized form of the FGL1 vector (FGL1-TM) by fusing a transmembrane sequence^[16]^. Next, the FGL1-TM vector were transferred together with the PD-L1 vector by lentiviral infection into HEK-293T, HepG2, A549, and MSC to establish stable cell lines overexpressing FGL1/PD-L1 (**Figure 2C**). Western blotting analysis (**Figure 2D**) and confocal images (**Figure 2E**) cooperatively confirmed FGL1 and PD-L1 were co-expressed and localized on the cell membrane.

**Figure 2.**
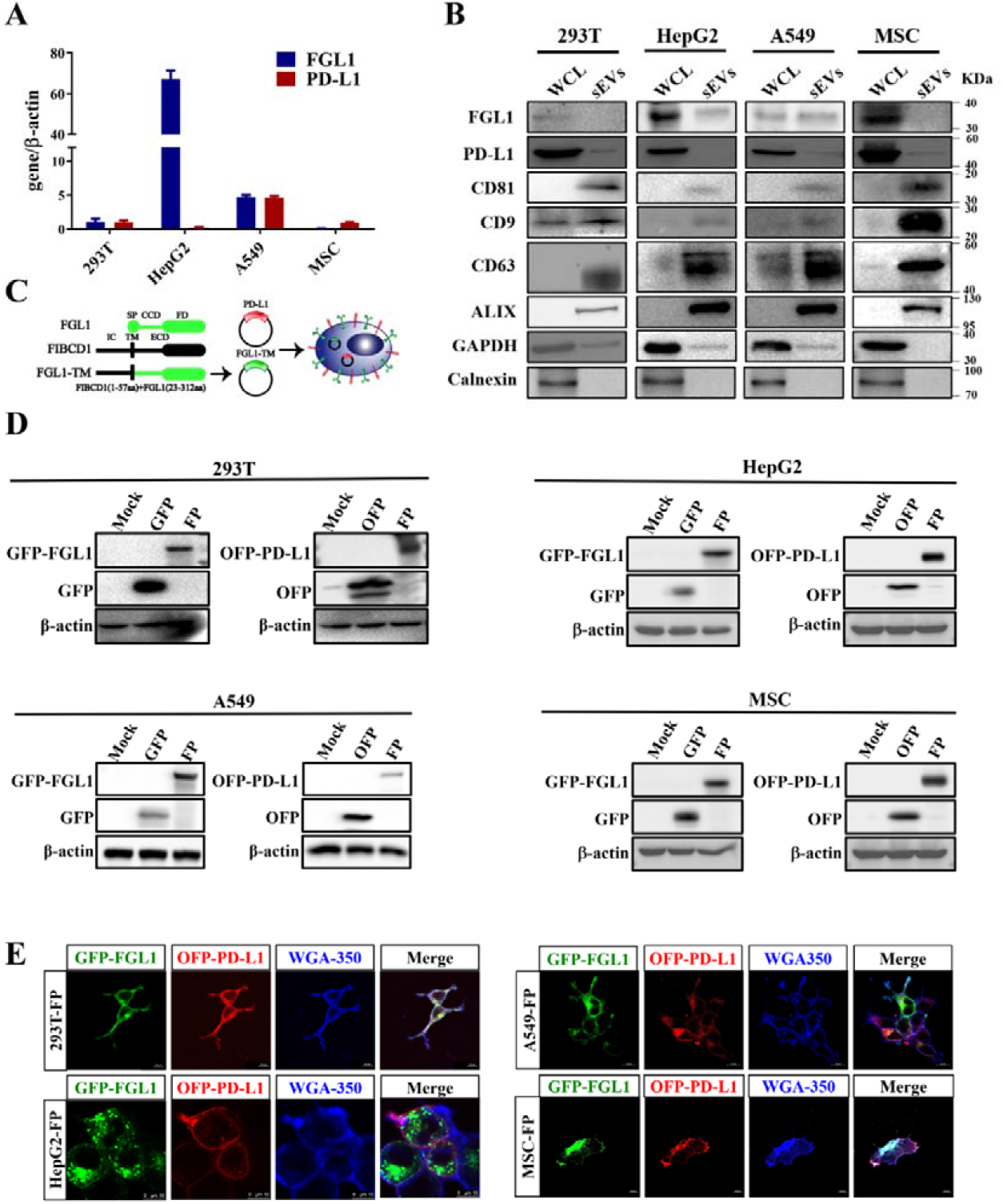
The deficiency of natural exosomes to express FGL1/PD-L1 simultaneously determines the necessity of artificial exogenous overexpression. A) The expression levels of FGL1 and PD-L1 in four different natural cell lines were detected by quantitative PCR, *n* = 3. B) Western blotting for FGL1, PD-L1, CD81, CD9, CD63, ALIX, GAPDH and calnexin in the whole cell lysate (WCL) and purified sEVs from HEK-293T, HepG2, A549, and MSC cells. C) Schematic diagram of FGL1 and PD-L1 plasmid construction. The modified FGL1 constructs contains an artificial transmembrane domain (FGL1-TM), the coil-coil domain (CCD) and fibrinogen domain (FD) of FGL1 (23-312aa) was fused with the FIBCD intracellular domain (IC) and transmembrane (TM) (1-57aa). SP represents signal peptide; ECD represents the extracellular domain. PD-L1 plasmid has the transmembrane structure of TM. D) HEK-293T, HepG2, A549 and MSC cells stably expressing FGL1/PD-L1 were lysed by RIPA lysis buffer, and the expression of FGL1 and PD-L1 was detected by Western blot. E) The expression of FGL1/PD-L1 on HEK-293T, HepG2, A549 and MSC cell membranes were observed by confocal microscope. Scale bar: 10 *μ*m.

Next, we prepared and purified dual-targeting FGL1/PD-L1 sEVs (FP sEVs) derived from HEK-293T-FP, HepG2-FP, A549-FP, and MSC-FP cells by differential centrifugation to test if overexpressing FGL1 and PD-L1 in cells can induce the enrichment of FGL1 and PD-L1 on sEVs. Western blotting confirmed the co-existence of exogenous FGL1 and PD-L1 in the sEVs derived from HEK-293T, HepG2, A549, and MSC cell lines, indicated by the exosomal markers CD63, CD9, and Alix in isolated vesicles, which were much higher in sEVs derived from bioengineering cells (FP sEVs) than unmodified cells (sEVs) (**Figure 3A**). Interestingly, MSC-FP sEVs expressed much more FGL1/PD-L1 than FP sEVs derived from other cell lines. The transmission electron microscopy (TEM) images showed that sEVs derived from different cell lines were round-shaped and membrane-bound (**Figure 3B**). Dynamic light scattering (DLS) analysis further verified similar size and stability of all types of sEVs, with an average diameter of 110 nm and average zeta potential of −30 mV (**Figures 3C-D**). Confocal microscopy further verified the co-presence of FGL1-GFP and PD-L1-OFP in the MSCs sEVs (**Figure 3E**). In additon, *in vitro* binding assay showed that OFP and GFP antibody simultaneously specifically pulled down FGL1-GFP and PD-L1-OFP-expressed exosomes, suggesting genetically engineered FGL1 and PD-L1 were localized on the surface of sEVs (**Figure 3F**).

**Figure 3.**
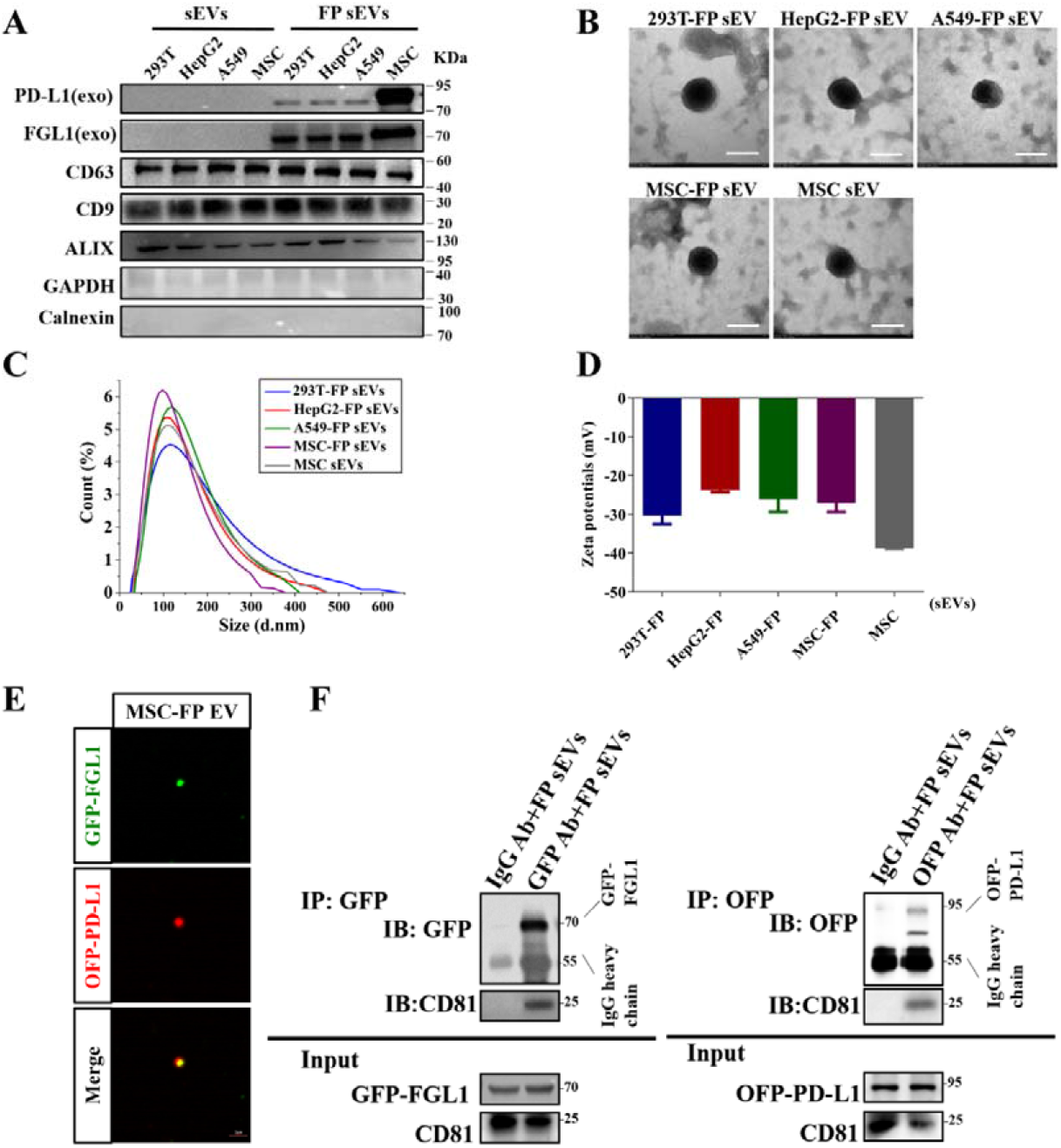
Characterization of FP sEVs and engineering MSC-FP can secrete modified exosomes delivering more FGL1/PD-L1. A) Western blotting for FGL1, PD-L1, CD9, CD63, ALIX, GAPDH and calnexin in the sEVs from HEK-293T, HepG2, A549, and MSC cells and overexpressed-FGL1/PD-L1 of these cell lines. B-D) The TEM images (B), size distribution (C), and the Zeta potential (D) of purified sEVs from HEK-293T-FP, HepG2-FP, A549-FP, MSC-FP and MSC cells, *n* = 3. E) Confocal images show the existence of FGL1-GFP/PD-L1-OFP on one sEV, scale bar: 2 *μ*m. F) FGL1/PD-L1 sEVs were incubated with GFP or OFP antibody followed with protein A/G Agarose pull-down. FGL1/PD-L1 and CD81 were detected by Western blotting. One percent of FGL1/PD-L1-expressed sEVs served as input.

As miRNAs derived from MSC sEVs are reported to maintain and regulate immune function^[17]^, we performed miRNA sequencing on HEK-293T and MSC-derived sEVs to verify if miRNAs from MSC sEVs have significant immunomodulatory potential. Using 4,872 Immunologic Signature Gene Sets provided by the GSEA website, miRNAs targeting these immune genes were mined from the original sequencing data and presented in the form of a heat map (**Figure S1A**). An extensive literature survey revealed that 9 out of the top 20 up-regulated and 2 out of top 20 down-regulated miRNAs have been proven to play a role in negative immune regulation (**Table S1**), of which we further validated three top-scored upgraded miRNAs (miR-125b-5p, let-7b-5p, and miR-21-5p) in MSC sEVs than HEK-293T sEVs by quantitative real-time PCR **(Figure S1B)**. In line with sequencing results, MSC sEVs showed stronger inhibition on PBMCs and T cells proliferation than HEK-293T sEVs through an *in vitro* CFSE cell proliferation assay (**Figure S1C**). Similar to MSC sEVs, MSC-FP sEVs also contained the same nine upregulated and two downregulated miRNAs of negative immunoregulation function when compared with HEK-293T sEVs (**Figure S2 and Table S1**), indicating that our genetic modification does not affect major immune-modulatory miRNA signatures in sE<u>V</u>s. These data suggested that MSCs were the optimal choice for bioengineering dual-targeting FGL1/PD-L1 sEVs due to their abundant expression of target proteins and potent immune regulation capacity. In addition, we constructed MSC cell lines overexpressing single target FGL1 or PD-L1 and GFP/OFP vectors (**Figure S3**), single-targeting PD-L1 and FGL1 sEVs were also prepared from MSCs and subjected to DLS analysis. There were no statistically significant differences in size, distribution pattern, and zeta potentials between single target sEVs and dual-targeting FGL1/PD-L1 sEVs (FP sEVs) (**Figure S4**).

### 2.3 *In vitro* inhibition of T cell proliferation by FGL1/PD-L1 interacting with their receptors, LAG3 and PD-1

To investigate whether FGL1 or PD-L1 sEVs were able to effectively bind to their target molecules, we conducted the following experiments. First, PD-L1 or FGL1 sEVs were incubated with OFP-LAG-3- and GFP-PD-1-expressing HEK-293T cells respectively. Confocal images revealed a distinct colorization of FGL1-GFP/LAG-3-OFP and PD-L1-OFP/PD-1-GFP on the membranes of HEK-293T (**Figure 4A**). As known, FGL1/LAG3 and PD-L1/PD-1 were two immune-negative regulatory pathways that played a major role in inhibiting T cell activation, so it was necessary to explore whether modified dual-targeting FGL1/PD-L1 sEVs (FP sEVs) could interact with LAG3/PD-1 expressing T cells. Consistent with the results of the heart transplant model (**Figure 1A**), the mRNA levels showed that LAG-3 and PD-1 expression increased when Jurkat cells were stimulated by PMA/ Ionomycin (PI) (**Figure S5**). FP sEVs were subsequently incubated with PI-stimulated Jurkat T cells. Confocal microscopy revealed a distinct co-localization of FP sEVs on the surface of activated Jurkat cells (**Figure 4B**). This indicates that FGL1/PD-L1 sEVs are able to bind specifically to LAG-3/PD-1 receptors.

**Figure 4.**
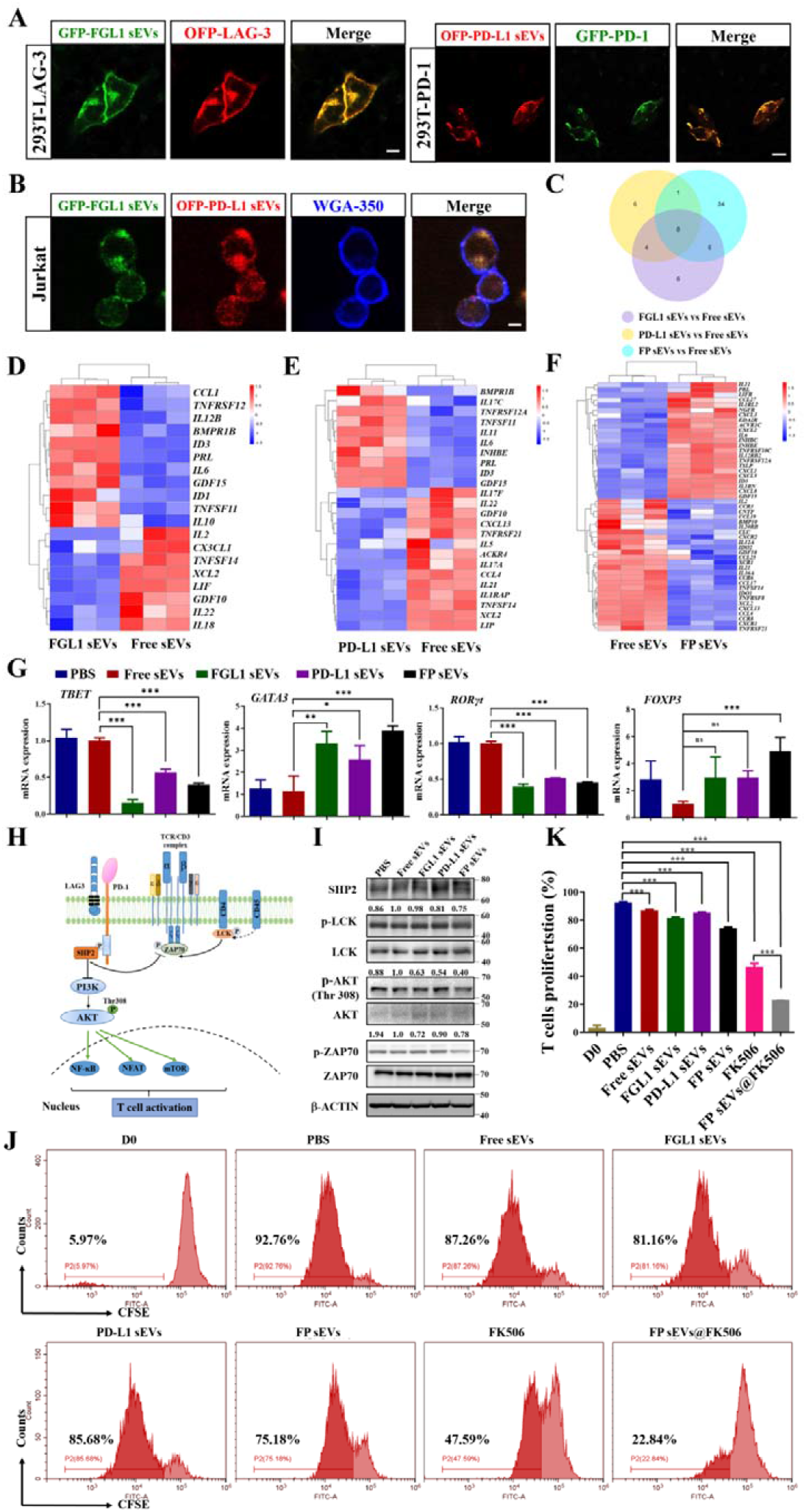
FGL1/PD-L1 sEVs could inhibit T cell proliferation and function. A-B) FGL1, PD-L1, and FGL1/PD-L1 sEVs interacted with the membrane of LAG-3-OFP expressing HEK-293T cells, PD-1-GFP of HEK-293T cells (A), and PI-stimulated Jurkat cells (B) for 30 min, respectively. Scale bar: 10 *μ*m. C-F) Transcript abundance was measured via Illumina RNA-seq analysis of a VENN diagram (C) and heatmap. D-F) of cytokine-related gene expression in FGL1 sEVs vs Free sEVs (D), PD-L1 sEVs vs Free sEVs (E) and FP sEVs vs Free sEVs (F). G) Quantitative PCR to verify the expression of *TBET*, *GATA3*, *RORγt*, and *FOXP3* in PBMCs after treating with different sEVs groups, *n* = 3. Error bar, mean ± SEM. *P*-values are calculated using one-way ANOVA by Tukey post-hoc test. H) PD-1/LAG3 involved in the classic TCR signaling pathway. I) Western blotting for TCR pathway protein expressions in Jurkat cells after treating with different sEVs groups. J) Different groups inhibited the proliferation of CD3^+^ T cells at 7 days shown using CFSE staining. CD3^+^ T cells were stimulated with CD3 and IL-2. After 3 days, sEVs (50 μg mL^-1^) were added to the cells for 7 days. CFSE staining was analyzed by flow cytometry. K) The quantitative analysis of T cell proliferation after culture with different groups of sEVs, *n* = 3. Error bar, mean ± SEM. *P*-values are calculated using one-way ANOVA by Tukey post-hoc test. \**P* < 0.05, \*\**P* < 0.01, \*\*\**P* < 0.001.

As overactive T cells are negatively regulated through the co-inhibitory pathway comprising FGL1/LAG-3 and PD-L1/PD-1, we explored the functional role of FP sEVs on the inhibition of T cell activation. First, healthy human PBMCs were extracted and co-incubated with Free sEVs (from control vector infected MSC), FGL1 sEVs, PD-L1 sEVs, and FP sEVs respectively. The mRNA resulting from each treatment was extracted and subjected to RNA-seq analysis after 48 h. It was immediately apparent that the concentration and number of regulatory cytokines were different between single target FGL1 or PD-L1 sEVs, and double target FP sEVs as compared to Free sEVs. However, FP sEVs had 9 and 14 overlapping cytokines as compared to the FGL1 sEVs and PD-L1 sEVs treatment groups respectively. In addition, the specific changes of 34 cytokines in the FP sEVs treatment group may be the reason for stronger efficacy observed than that of any single target treatment group (**Figure 4C**). For example, FGL1 sEVs decreased *IL-2*, *IL-22*, and *IL-18*, and increased *IL-10* as compared with the Free sEVs group (**Figure 4D)**. It is believed that the decrease of *IL-2* can activate Treg cells, *IL-18* and *IL-22* are associated with the differentiation of CD4^+^ T cells (**Table S2**)^[18]^, while *IL-10* can alleviate immune rejection accompanied by rebuilding immune tolerance^[19]^. *IL-5*, *IL-17A*, *IL-17F*, and *IL-21* decreased in PD-L1 sEVs compared with Free sEVs (**Figure 4E)**. According to authoritative literature reports, *IL-5* can mediate allogeneic immune rejection^[20]^, *IL-17A* and *IL-17F* have been shown to promote the differentiation of CD4^+^ T cells^[18b]^, while *IL-21* can inhibit the proliferation of Treg cells (**Table S2**)^[21]^. In comparison to the Free sEVs group, the changes of cytokines in the FP sEVs group were more prominent (**Figure 4F)**. In conclusion, the co-expression of the FGL1/PD-L1 can affect the proliferation and differentiation of immune cells to some extent by affecting more immune cytokines.

In addition to RNA-sequencing, we wanted to analyze the effect of FP sEVs on the different CD4^+^ T-helper-cell (Th1/Th2/Th17) subsets and regulatory T-cells (Tregs), we first detected the mRNA levels of specific transcription factors in Th1, Th2, Th17, and Treg cells (**Figure 4G**). We found that *TBET* and *RORγt* transcription factors were down-regulated, indicating that our dual-targeting sEVs were able to suppress the differentiation of Th1 and Th17, thus inhibiting the function of T cell proliferation and differentiation. They also affected a concomitant increase in *FOXP3*, indicating an ability to increase the proportion of Treg cells. The sEVs were also able to increase the expression of *GATA3*, which is suggestive of promotion in Th2 differentiation. According to previous reports, down-regulation of the Th1/Th2 ratio promotes stability of the immune environment after organ transplantation. On the basis of the above results, we further investigated simultaneously the phenotypes of effector Th-cells that express Th1 (IFN-γ) and Th17 (IL-17A) cytokines as well as Tregs. There was a reduced percentage of Th1 and Th17 cells along with increased Treg cells **(Figure S6)**. The results were consistent with those of the above variable transcription factors. All the above evidence demonstrated FP sEVs could suppress T cell function to inhibit immune rejection via inducing an inhibitory T helper cell differentiation type.

In order to clarify how our artificially modified sEVs affected downstream signaling in the T cell receptor (TCR) pathway, Jurkat cells were lysed in order to observe related proteins after 48 h incubation with different sEVs groups. PD-1/LAG3 involved in TCR pathway is clearly shown in **Figure 4H.** The results show that p-LCK, p-AKT (Thr308), p-ZAP70 were down-regulated, thus inhibiting T cell proliferation and activation. SHP2 is a signaling protein recruited by the synergistic action of LAG-3 and PD-1. Its role is to inhibit rapid protein phosphorylation occurring at T cell downstream signaling sites, thereby weakening T cell function (**Figure 4I**). Additionally, FP sEVs showed a stronger effect on elevating SHP2 while reducing p-LCK and p-AKT (Thr308) expression compared with FGL1 sEVs or PD-L1 sEVs. Therefore, the modified sEVs could inhibit T cell proliferation and T cell function via suppressing immune cytokines, T cell differentiation, and TCR downstream pathway proteins.

Lastly, CFSE-labeled CD3^+^ T cells were cultured with Free sEVs, FGL1 sEVs, PD-L1 sEVs, and FP sEVs respectively for 7 days. The following flow cytometry analysis showed that FGL1, PD-L1, and FP sEVs significantly inhibited the proliferation of CD3^+^ T cells by 11.6%, 7.08%, and 17.58% respectively, suggesting that FP sEVs had a stronger immunosuppressive effect than that of FGL1 or PD-L1 single-targeted sEVs (**Figure 4J**). This was supported by a corresponding quantitative analysis of CFSE results (**Figure 4K**).

FK506 is the most widely used immunosuppressive agent in tissue and organ transplantation rejection and acts by preventing Ca^2+^-calcineurin-NFAT signaling, the master regulator of T cell proliferation and activation^[22]^. To investigate the efficacy of FGL1/PD-L1 sEVs as a synergistic, targeted drug delivery system of FK506 to effector T cells, we prepared coated vesicles (FP sEVs@FK506) using an electric transformer. The average drug loading percentage was 23.58% (**Figure S7**). As compared to FK506 and FP sEVs administered alone, FP sEVs@FK506 achieved the highest inhibition rate of 69.92% on T cell proliferation (**Figure 4J-K)**, suggesting that FP sEVs@FK506 greatly enhanced the inhibition function of FK506.

### 2.4 FGL1/PD-L1 sEVs prolonged graft survival time and induced immune tolerance in a heart graft model

We performed verification experiments in a mouse cardiac allograft model, an excellent tool to study immunological mechanisms^[23]^. The hearts of BALB/c mice were transplanted to the cervices of C57BL/6 mice. We first infected mouse FGL1/PD-L1 on MSC cells to obtain stable cell lines (**Figure S8**) and then extracted enough sEVs for drug administration to further assess whether FP sEVs showed better inhibitory effect in transplant rejection. The recipient mice were then grouped into seven clusters containing saline, Free sEVs, PD-L1 sEVs, FGL1 sEVs, FP sEVs, FK506, and FP sEVs@FK506 (**Figure 5A**). Bioluminescent images of FP sEVs taken *in vivo* confirmed a marked increase in accumulation on cervical graft-heart sites as compared to both sEVs (from natural MSCs) and Free sEVs (from control vector infected MSCs) (**Figure S9A**). There was no significant difference in body weight change among each group (**Figure 5B**). Simultaneously, toxicological experiments were also performed, including a histopathological assay and blood cell counts, which proved that our sEVs did not produce other obvious side effects elsewhere in the mice (**Figure S9B-C)**. In the course of the experiment, we were also surprised to find that multiple injections of FK506 caused rough, matte, depilation of mouse hair, and elevated creatinine compared to FP sEVs group. It was reflective that FP sEVs@FK506 could reduce some of the toxic side effects associated with FK506 alone (**Figure S10)**. Among these groups, FP sEVs showed notably prolonged heart graft survival **(Figure 5C)**, and a declining trend of CD8^+^ T cells was observed in spleens and grafted hearts (Figure 5D-E). In contrast to this, CD4^+^ CD25^+^ Foxp3^+^ regulatory T cells (Treg cells) were more prevalent in the superficial cervical lymph nodes of heart-transplanted mice that were injected with FP sEVs **(Figure 5F)**. Similarly, FP sEVs could cooperate with FK506 (FP sEVs@FK506) showed the greatest propensity to reduce the number of CD8^+^ T cells in spleens and transplanted hearts (Figure 5D-E), and caused an increase in CD4^+^CD25^+^Foxp3^+^ Treg cells (**Figure 5F**). Remarkably, FP sEVs and FP sEVs@FK506 precluded immune rejection via decreasing levels of inflammatory cells and infiltrated CD3^+^ T cells (**Figure 5G**). We further detected cytokines of Granzyme B, IL-12/IL23p40, TNF-*α*, including TGF-*β*1 from serum as the presence of cytokines is also reflective of the activation capacity of immune cells. FGL1 sEVs, PD-L1 sEVs, FP sEVs, and FP sEVs@FK506 treatment groups all displayed a prominent reduction of Granzyme B, IL-12/IL23p40, TNF-*α*, additionally with a noticeable rise in TGF-*β*1 **(Figure 5H)**. Overall, this supports the conclusion that FP sEVs exhibit a synergistic immunosuppressive function with FK506 to prevent graft rejection and reestablish immune tolerance.

**Figure 5.**
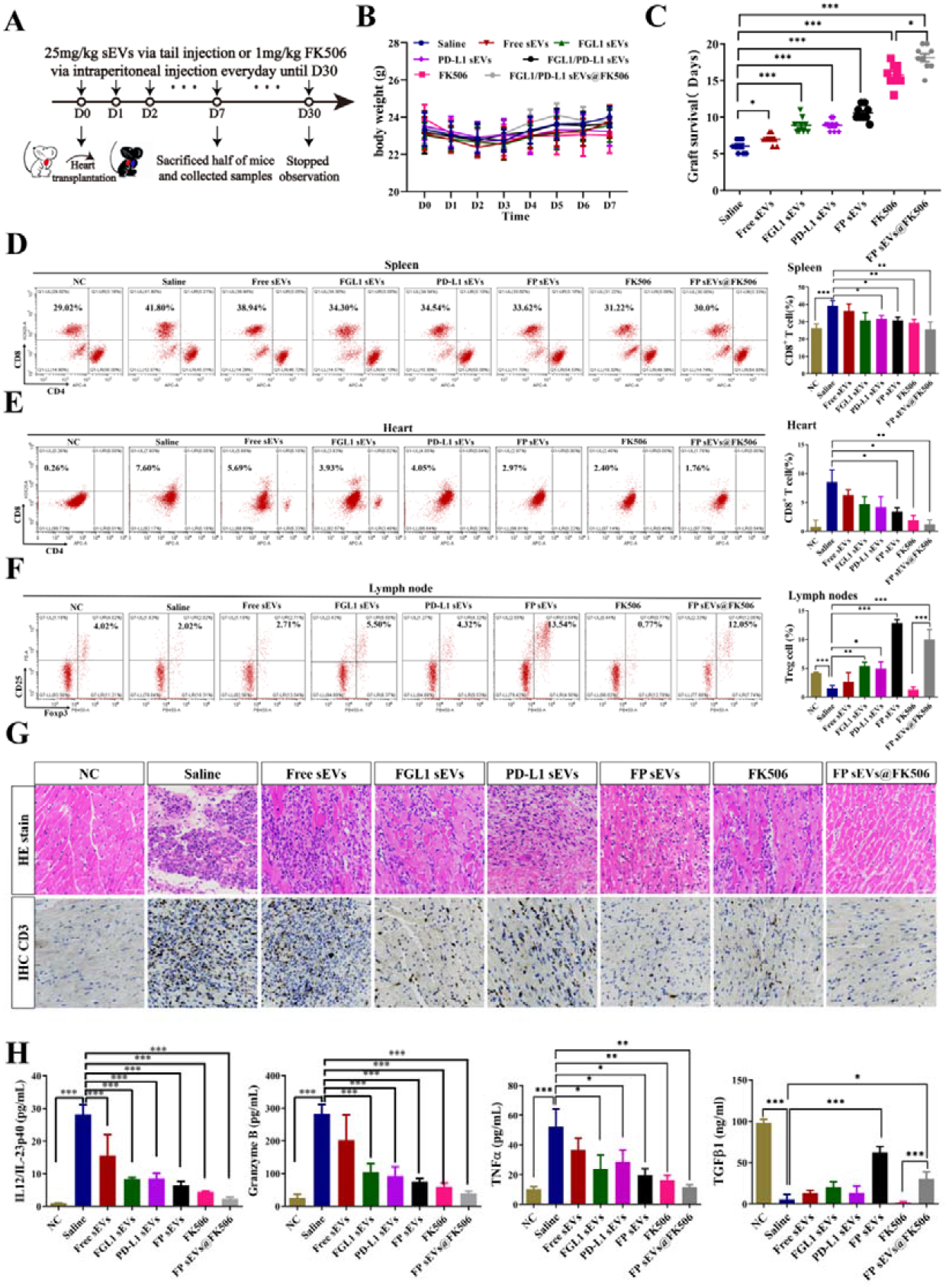
FGL1/PD-L1 sEVs promotes prolonged graft survival time and immune tolerance in a heart graft model. (A) Images of the mode of administration of every group treatment. (B) The body weight of heat-graft mice in diverse groups as illustrated, *n* = 9. (C) Graft survival curves of cardiac allografts treated with different groups (Saline, Free sEVs, PD-L1 sEVs, FGL1 sEVs, FGL1/PD-L1 sEVs, FK506, FGL1/PD-L1 sEVs@FK506), *n* = 9. Error bar, mean ± SEM. *P*-values are calculated using one-way ANOVA by Tukey post-hoc test. D-E) Variation of CD8+ T cells in spleens (D) and hearts (E) were measured by flow cytometry separately, *n* = 5. F) Characteristic flow cytometry charts reveal changes of CD4^+^CD25^+^Foxp3^+^ Treg cells in the superficial cervical lymph nodes of heart-transplanted mice, *n* = 3. G) Hematoxylin-eosin (HE) staining and CD3-immunohistochemistry (IHC) show inflammation changes and quantity variance of infiltrated CD3^+^ T cells in grafted-heart, respectively. Scar bar: 50 *μ*m. H) Secretion of cytokines for IL-12/IL23p40, Granzyme B, TNF-*α* and TGF-*β*1 in serum by ELISA, *n* = 5. Error bar, mean ±SEM. *P*-values are calculated using one-way ANOVA by Tukey post-hoc test. \**P* < 0.05, \*\**P* < 0.01, \*\*\**P* < 0.001.

## 3. Discussion and Conclusion

In summary, we established and designed MSC-derived dual-targeting FGL1/PD-L1 sEVs for inhibiting immunological rejection. This dual-targeting drug delivery system also exhibits a strong ability to carry low doses of FK506 to LAG-3/PD-1 expressing effector T cells, thus inhibiting T cell activation and proliferation, and inducing Tregs in organ recipient mice. Our study provides an experimental basis for a novel intervention strategy that leverages the function of target delivery sEVs to synergistically enhance two immunosuppression axes and reestablish immune tolerance, ultimately promoting organ acceptance **(Figure 6)**.

**Figure 6.**
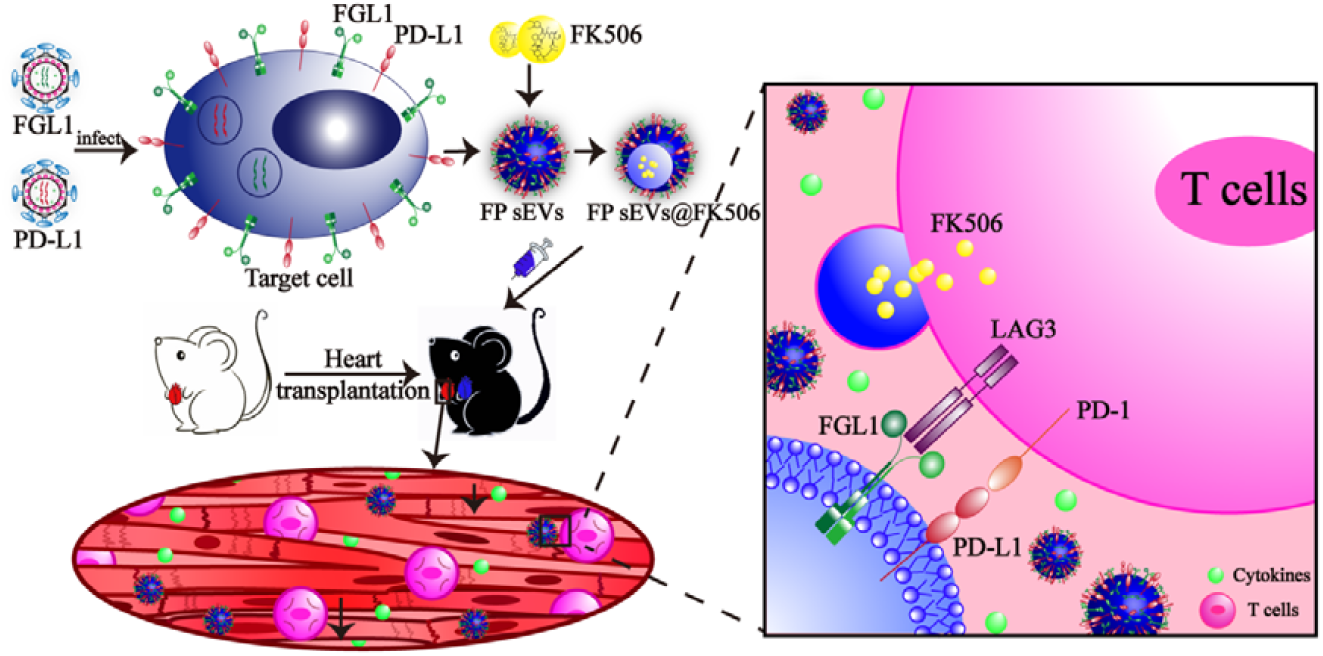
A schematic model showing FP sEVs@FK506 suppress T cell activation and thus inhibit cardiac allograft rejection. FGL1 and PD-L1 packed with lentivirus successively infected target cells and thus were stably expressed upon cell surfaces. sEVs secreted by engineered cells stably expressed FGL1/PD-L1. At the same time, FK506, a classic immunosuppressant, was introduced into sEVs by electroporation. We translated the effects of FGL1/PD-L1 sEVs@FK506 onto a mouse heart-graft model to test their immunosuppressive function *in vivo*. FGL1/PD-L1 sEVs@FK506 administered by tail vein injection migrated to the grafted-hearts, causing a marked reduction in T cell activation and cytokine secretion. This was attributed to the blockage of the PD-L1/PD-1 and FGL1/LAG-3 axes, as well as the inhibitory effects of FK506 in immunological ejection.

LAG-3 is reported to be synergistic with PD-1 inhibitory effects on T cell signaling during immune evasion in cancer^[24]^. Dual antibody blockade or genetic knockout of LAG-3 and PD-1 significantly enhanced T effector function and delayed tumor growth. In this study, LAG-3 and PD-1 were detected simultaneously up-regulated expression under an organ transplant rejection environment so as to maintain immune tolerance, providing a solid foundation for the concurrent construction of double-targeting immunosuppressants. Membrane-based nanovesicles including erythrocytes, platelets, or nanovesicles obtained by the grinding and centrifugation of cell membranes have enormous potential to behave as drug delivery vehicles owing to their small size and biocompatibility^[25]^. However, there are several problems associated with their preparation, including membrane incompleteness, turnover of inner and outer membranes, especially high immunogenicity. As a natural membrane delivery system, sEVs present themselves as an excellent alternative biological vector. However, whether two ligand proteins can be simultaneously highly expressed on the membrane of sEVs by genetic engineering has not been reported before. Herein, we presented the first report on immune checkpoints dual-targeting sEVs and their application in organ transplantation. Furthermore, we determined that MSC-derived sEVs prior to that from other cell sources can inherit the properties of immune modulation and tolerance of MSCs themselves, and will therefore be a promising biomaterial for immunotherapies.

We modified sEVs to simultaneously carry two targets exogenously, but it is unclear whether such a manipulation affects sEVs quality and function. Indeed, the difficulty in controlling the quality and function of modified sEVs remains a substantial obstacle in their development as a novel therapeutic strategy. We therefore compared sEVs from a diverse range of cell sources and also analyzed the contents of engineered sEVs, which led to the conclusion that the variation in sEVs contents should be closely monitored in order to ensure a high level of quality control during the modification process.

Tregs play an important role in establishing immune tolerance, preventing an excessive immune response and eliminating autoimmunity. Tregs have been widely studied as a cell therapy in the treatment of GvHD and the limiting of graft rejection. However, previous studies have reported that Foxp3 in Treg cells are less NFAT-independent, so two of the most commonly used immunosuppressants of FK506 and Cyclosporin A, the inhibitors of NFAT signaling pathway, cannot effectively reconstruct Treg cell subsets in patients which display organ transplantation rejection^[22b, 26]^. Excitingly, we found that FP sEVs@FK506 additionally induce regulatory T cells in the recipient surgical lymph nodes, indicating the power of combining low-dose FK506 with FGL1/PD-L1 on sEVs as immunosuppressants to promote allograft acceptance. This underscores the remarkable function of immune checkpoints in excessive immunity. In the future, bioengineering sEVs carrying multiple disease-mediating receptors or cross-talking signaling cascades may become a common therapeutic strategy.

## 4. Experimental Section

### 4.1 Clinical samples of kidney transplant patients

Nineteen kidney transplant recipients were enrolled in this study. They were divided into three groups: stable group (*n* = 7), antibody-mediated rejection (ABMR) group (*n* = 6), and T cell-mediated rejection (TCMR) group (*n* = 6). Approximately 20 mL of heparinized peripheral blood was obtained after allograft transplantation from patients. All patients gave informed consent for this study, which was approved by the Organ Transplant Center at the First Affiliated Hospital of Sun Yat-sen University (Clinical Trials. gov NO. 2019-456). Refer to previously published articles for the demographic clinical characteristics of the study in kidney transplant patients^[22b]^. Information of patients involved in the experiments are listed in the **Table S3**

### 4.2 Biochemicals and Antibodies

Puromycin was purchased from Sigma-Aldrich. GAPDH, *β*-actin, GFP, OFP, SHP2, p-LCK, LCK, p-AKT (Thr308), AKT, p-ZAP70, and ZAP70 antibodies for western blot were purchased from Abmart. FGL1 antibodies were purchased from Santa Cruz Biotechnology Inc. Antibodies, including Human PD-L1 and PD-1 were purchased from Invitrogen. Human LAG-3 and Na^+^K^+^ATPase antibodies were purchased from Cell Signaling Technology and Santa Cruz Biotechnology Inc. respectively. Marker antibodies for exosomes, including anti-CD9, anti-CD63, and anti-ALIX, were purchased from Santa Cruz Biotechnology Inc, and anti-CD81 from System Biosciences. Wheat Germ Agglutinin (WGA) Alexa Fluor 488 and 350 dyes were purchased from Thermo Scientific. Ficoll Paque Plus used for isolating peripheral blood mononuclear cells (PBMC) cells was purchased from GE Healthcare. Staining antibodies, including CD3, CD4, CD8, CD25, and Foxp3 for FACS analysis were purchased from Biolegend Inc.

### 4.3 Cell lines and Cell cultures

HEK-293T cells (human embryonic kidney cell lines), Jurkat cells (human acute T cell leukemia cell lines), HepG2 cells (human liver cancer cell lines), and A549 cells (human lung cancer cell lines) were purchased from American Type Culture Collection (ATCC). Cells were cultured in RPMI 1640 or DMEM supplemented with 10% fetal bovine serum (Gibco), 100 units mL^-1^ penicillin, and 100 μg mL^-1^ streptomycin, and were incubated at 37 °C in a 5% CO_2_ atmosphere.

### 4.4 Plasmids and stable cell lines

Plasmids from human (pLV-puro-TM-FGL1-GFPSpark), (pLV-puro-PD-L1-OFPSpark), (pLV-puro-PD-1-GFPSpark), and (pLV-puro-LAG-3-OFPSpark) and plasmids from mouse (pLV-puro-TM-FGL1-GFPSpark), and pLV-puro-mPD-L1-OFPSpark) were purchased from Sino Biological Inc. For stable cell lines, HEK-293T cells were transfected with packaging, envelope, and target plasmids using Lipo 3000 (ThermoFisher, Waltham, USA). Fluid change was performed at 12 h and at 24, 48, and 72 h post fluid change. Lentivirus-containing supernatant was collected, filtered with a 0.45 *μ*m filter membrane, and stored at −80°C. HEK-293T, HepG2 cells, and A549 cells were infected with lentivirus and selected with puromycin (2 μg mL^-1^) to obtain stable expressing target cells.

### 4.5 Generation and purification of sEVs

Cells and genetically engineered cells were grown in 15 mm culture dishes and allowed to proliferate. When cells reached 80% confluence, the culture medium was replaced with a similar volume of DMEM supplemented with 0.5% exosome-free FBS [the supernatants were centrifuged at 100,000 × *g* for 12 h (Beckman Coulter, Optima L-100XP) using normal FBS]. After incubating for 36–48 h, exosomes were extracted using traditional gradient centrifugation methods. To prevent exosome degradation, all centrifugation steps were performed at 4 °C. In brief, culture supernatants were centrifuged at 500 × *g* for 10 min, 2000 × *g* for 20 min and 10,000 × *g* for 40 min successively to remove dead cells, cell debris and other cell secretions (Beckman Coulter, Allegra X-30R). The supernatants were then centrifuged at 100,000 × *g* for 90 min (Beckman Coulter, Optima L-100XP) to obtain the exosomes. The pelleted exosomes were resuspended in 50–150 μL precooled PBS and stored at −80 °C immediately for the following experiments. It should be noted that in general, pellets were suspended in RIPA lysis buffer (Thermo Scientific) only for western blot.

### 4.6 Extraction and culture of umbilical cord mesenchymal stem cells (UC-MSCs)

Fresh umbilical cord of approximately 10 cm was obtained after cesarean section, the remaining blood of the umbilical cord was washed thoroughly with normal saline, and then cut into small segments of 2–3 cm, and then rinsed again. The umbilical cord was cut vertically, then one umbilical vein and two umbilical arteries were removed, and Huatong glue was extracted. Ophthalmic scissors were used to cut the Huatong glue into small tissue blocks of 1 mm, which were then transferred to cell culture bottles. DMEM/F12 was added to culture medium containing 10% FBS, 1× penicillin/streptomycin, then cultured in an incubator with 5% CO_2_ and at 37°C for static cultures. Cell growth was observed under an inverted microscope every day. The liquid was changed for the first time after 1 week, and once every 3–4 days thereafter. When the cells were 80% ~ 90% long, trypsin/EDTA digestion solution was used for digestion and passage.

### 4.7 Western blot

Cell lysates and exosomes including purified membrane vesicles were separated by SDS-PAGE and were then transferred to polyvinylidence fluoride membranes (Millipore, Darmstadt, Germany). The membrane was sealed with 5% non-fat milk for more than 1 h at room temperature and incubated with the desired primary antibodies overnight at 4 °C. Post incubation with HRP-conjugated secondary antibodies was performed for 1 h at room temperature, then detection was carried out using enhanced chemiluminescence reagent (ECL) (Protein Tech, China).

### 4.8 Characterization of sEVs

The size and zeta potential of exosomes in PBS were determined using a NanoBrook 90Plus PALS (Brookhaven instruments). Data recorded are the average of three measurements. A transmission electron microscopy (TEM, hc-1, Hitachi) at 80 kV was used to observe the morphology of the exosomes. 10 μL of purified exosomes were suspended in PBS and placed on formvar-carbon-coated copper grids. After 10 min, the residual liquid was removed from the grid edge with filter paper. Exosomes on the grids were then stained with 2% uranyl acetate for 10 min, washed with deionized water 3 times, each time for 10 min, then air-dried.

### 4.9 sEVs cell binding assay

For HEK-293T-LAG-3-OFP and HEK-293T-PD-1-GFP cells, cells were seeded respectively in confocal dishes, incubated with FP sEVs for 30 minutes the next day, then membranes were stained with WGA 350 for 15 min and observed with a confocal microscope. Jurkat cells were pre-stained with WGA 350 for 15 min and incubated with FP sEVs for 30 min, then spun onto glass slides and observed by confocal microscopy (Zeiss, LSM880).

### 4.10 FK506 loading

The mixtures containing 200 μg FK506 and 1 mg sEVs diluted in PBS were added to 0.4 cm electroporation cuvettes and were subjected to electroporation at 300 V and 150 μF using a Bio-Rad Gene Pulser Xcell Electroporation System. For membrane recovery, samples in electroporation cuvettes were incubated on ice for 30 min and rinsed gently with PBS to gain suspension. Following centrifugation at 12,000 × *g* for 10 min, the resulting pellets were washed with cold PBS 3 times and resuspended in PBS for further application.

### 4.11 Isolation of PBMC and T cells from human peripheral blood

Peripheral blood from normal healthy donors was collected into EDTA potassium vacuoles. PBMCs were isolated using Ficoll lymphocyte isolation medium. Next, CD3^+^ T lymphocytes were purified (> 98%) by negative selection with MojoSort^™^ Human CD3^+^ T cell Isolation Kit (Biolegend, USA). Cells were seeded in CD3-coated plates (clone OKT3; Biolegend) and cultured in medium (RPMI1640 with 10 % FBS, penicillin/streptomycin, and 2 ng mL^-1^ IL-2).

### 4.12 CFSE staining

A CFSE separation and tracking kit (Biolegend, USA) was used to label PBMCs as a CFSE working solution (5 μM). T cells were incubated at 37 °C for 20 min, and then quenched and stained with culture medium on 0, 3, 5, and 7 days respectively. Cells were collected and subjected to FACS (Cytoflex, Beckman, USA) using the Cell Quest software (CytExpert, USA).

### 4.13 Intracellular Flow Cytometry Staining

For the percentage analysis of Th1, Th17 and Treg cells, the superficial cervical lymph nodes on the surgical side of recipient mice were harvested and a single-cell suspension was prepared as described above. Cells first treated with BV421-labeled anti-human CD4, then fixed by fixing buffer for 20 min, permeated with Intracellular Staining Perm Wash Buffer (Biolegend), and then stained with PE anti-human IL-4, Alexa Fluor® 647 anti-human IL-17A antibody and APC/PC7 anti-human IFN-γ. Cells were collected and subjected to FACS (Cytoflex, Beckman, USA) using the Cell Quest software (CytExpert, USA).

### 4.14 Biological distribution of sEVs

Cy5.5-labeled sEVs, Free sEVs and FP sEVs were injected into BALB/c mice as a 200 μL (2 μg μL^-1^) solution via the tail vein. After 2 h, accumulation of sEVs in various organs was observed using a NightOWLimaging system (LB983).

### 4.15 Mouse heart graft model

The use of laboratory animals and all animal experiments was reviewed and approved by the Animal Ethics Committee of the Zhongshan School of Medicine, Sun Yat-sen University, China. The approval number is SYSU-IACUC-2019-000332. Heart transplantations were carried out using male BALB/c mice as donors. Hearts were transplanted into the necks of male C57BL/6 mice (8-10 weeks old, weight >22 g). The doses and injection time of all 78 recipients were randomly divided into four equal groups: group 1 (negative control group, n = 8), group2 (injected with saline, n = 10), group 3 (Free sEVs 25 mg kg^-1^, n = 10), group 4 (FGL1 sEVs 25 mg kg^-1^, n = 10), group 5 (PD-L1 sEVs 25 mg kg^-1^, n = 10), group 6 (FK506 1.0 mg kg^-1^, n = 10), group 7 (FGL1/PD-L1 sEVs 25 mg kg^-1^, n = 10), and group 8 (FGL1/PD-L1 sEVs@FK506 25 mg kg^-1^, n = 10). The recipients were treated via tail vein injection every day, until 30 days after transplantation. Transplanted mice were then sacrificed at 7 days to dissect grafted-hearts, lymph nodes, and spleens, which were subjected to downstream analysis.

### 4.16 Histology and immunohistochemistry analysis

Seven days after heart graft surgery, the mouse recipients were sacrificed and tissue examples from the transplantation location were collected in 15 mL centrifugal tubes, and the entirety of samples were submerged with 4% paraformaldehyde for fixation. Next, the samples were embedded with paraffin and were sectioned (4 *μ*m thickness). Sections were stained with hematoxylin and eosin (H&E) using standard procedures. Inflammatory cell infiltration conditions were observed by fluorescence microscopy with 4x, 10x, 20x and 40x magnification. For the immunohistochemical staining, deparaffinized tissue sections were incubated with mouse monoclonal CD3 (PC3/188A: sc-20047, Santa Cruz Biotechnology), followed by visualization using an HRP/DAB detection IHC Kit. Sections were counterstained with Mayer’s hematoxylin. The infiltrated parts of lymphocytes were also photographed as above.

### 4.17 RNA isolation and qPCR analysis

The total RNA was collected and purified from cells or spleen tissue using TRIZOL reagent (TaKaRa, Tokyo, Japan) according to the manufacturer’s protocol. RNA concentrations were measured using NANODROP ONE (Thermo Fisher Scientific). RNA was reversely transcribed into complementary DNA (cDNA) using HiScript III RT SuperMix for qPCR (+gDNA wiper) (TransGen Biotech, China) with a PCR Instrument (BIO-RAD), and then quantified by qPCR using 2x SYBR Green qPCR Mix (TransGen Biotech, China) with LightCycler® 96 (Roche). Relative gene expression folding changes were identified with the 2^-ΔΔCt^ method. The qPCR primers for all the experiments are listed in **Table S4**.

### 4.18 RNA Sequencing

RNA extraction and qualification: PBMCs were extracted from healthy volunteers. sEVs from different treatment groups were co-incubated with PBMCs for 48 h, then RNA was extracted using TRIZOL reagent (TaKaRa, Tokyo, Japan). RNA degradation and contamination were monitored on 1% agarose gels. RNA purity and concentration were checked using a NanoPhotometer® spectrophotometer (IMPLEN, CA, USA) and Qubit® RNA Assay Kit in Qubit® 2.0 Flurometer (Life Technologies, CA, USA). RNA integrity was assessed using an RNA Nano 6000 Assay Kit from the Bioanalyzer 2100 system (Agilent Technologies, CA, USA). The RNA-seq library was then generated using the NEBNext® Ultra^™^ RNA Library Prep Kit for Illumina® (NEB, USA). Library quality was assessed using the Agilent Bioanalyzer 2100 system. Subsequent sequencing was performed using an Illumina Novaseq System (Illumina, USA). Library construction and sequencing were carried out at Berry Genomics. For data analysis, differential expression analysis of two conditions was performed using the DEGSeq R package. The *P* values were adjusted using the Benjamini & Hochberg method. A corrected *P* value of 0.05 and log 2 (fold change) of 1 were set as the threshold for significantly differential expression.

### 4.19 MiRNA Sequencing

RNA from EVs was extracted using an RNA miRNeasy Micro Kit (Qiagen, Germany). RNA quantity and purity were determined by a Nanodrop (Thermo Fisher Scientific Inc., USA) and an Agilent 4200 Tape station (Agilent, CA, USA). Library construction for the paired-end libraries was performed using a QIAseq miRNA library kit (Qaigen, Germany). Subsequent sequencing was performed on the Illumina Novaseq System (Illumina, USA). Library construction and sequencing were carried out at Shanghai Biochip Corporation. For data analysis, the reading of each miRNA-seq sample was compared with the existing sequence in the miRBase and the predicted result of the new miRNA in order to calculate the expression level count (counts per million) of the miRNA. The edgeR software package was used to analyze the expression between samples and screen out UMI counts with a *P* value of <0.05 and a fold change of >2.0.

### 4.20 Isolation of CD4^+^ and CD8^+^ T cells from mice spleens

Spleens of the mice were dissected and placed in 1.5 mL EP tubes. A single-cell suspension was obtained by passing the suspension through a 100 *μ*M cell sieve. The cells were centrifuged at 1500 × *g* for 5 min at 4 °C, then resuspended and centrifuged twice with RBC lysis buffer to obtain cell pellets. Next, CD4^+^ T and CD8^+^ T lymphocytes were purified (> 98%) by negative selection with MojoSort^™^ Mouse CD4^+^ T cell Isolation Kit and MojoSort^™^ Mouse CD8^+^ T cell Isolation Kit (Biolegend, USA).

### 4.21 Flow cytometric analysis

Seven days after heart transplantation, the superficial cervical lymph nodes on the surgical side, graft-heart, and spleens of the mice were dissected and placed in 1.5 mL EP tubes. The hearts were cut with scissors, and the spleens and lymph nodes were ground with a mortar. A single cell suspension was obtained by passing the suspension through a 70-mesh cell sieve. The cells were centrifuged at 1500 × *g* for 5 min at 4 °C, then resuspended and centrifuged twice with RBC lysis buffer to obtain cell pellets. These mononuclear cell suspensions were labeled with FITC-CD3, APC-conjugated CD4, BV510-conjugated CD8, PE-conjugated CD25 and Pacific Blue-conjugated Foxp3 monoclonal antibodies, then subjected to FACS (Cytoflex, Beckman, USA) using the Cell Quest software (CytExpert, USA).

### 4.22 Statistical Analysis

Three independent sample replicates were carried out for each experiment unless stated otherwise. The statistical significance between the two groups was measured using the unpaired Student’s t-test. All results are expressed as the mean ± SEM. Data analysis and processing was done using GraphPad Prism Ver 8.0 (GraphPad Software). One-way analysis of variance (ANOVA) or Student’s T-test was performed, and statistical significance was indicated (*, *p* < 0.05; **, *p* < 0.01; ***, *p* < 0.001).

## Acknowledgements

H-i T., Y.Y.W. and X.Y.L contributed equally to this work. B.J, F.C, and H.B.C. are corresponding authors, This research was supported by the National Natural Science Foundation of China (81702750, 81970145 and 82001698); Natural Science Foundation of Guangdong Province (2020A1515011465 and 2020A151501467); Science, Technology & Innovation Commission of Shenzhen Municipality (JCYJ20180307154700308, JCYJ20170818163844015, JCYJ20180307151420045, JCYJ20190807151609464, JCYJ20200109142605909 and JCYJ20210324120007020); Sun Yat-sen University (20ykzd17 and 20ykpy122); International Collaboration of Science and Technology of Guangdong Province (2020A0505100031); Guangdong Provincial Key Laboratory of Digestive Cancer Research (No. 2021B1212040006); The Social Development Foundation of Jiangsu Province (BE2018691) and Sigrid Jusélius foundation in Finalnd for funding the project.

## Supporting Information

### Supplement Figures

**Figure S1.**
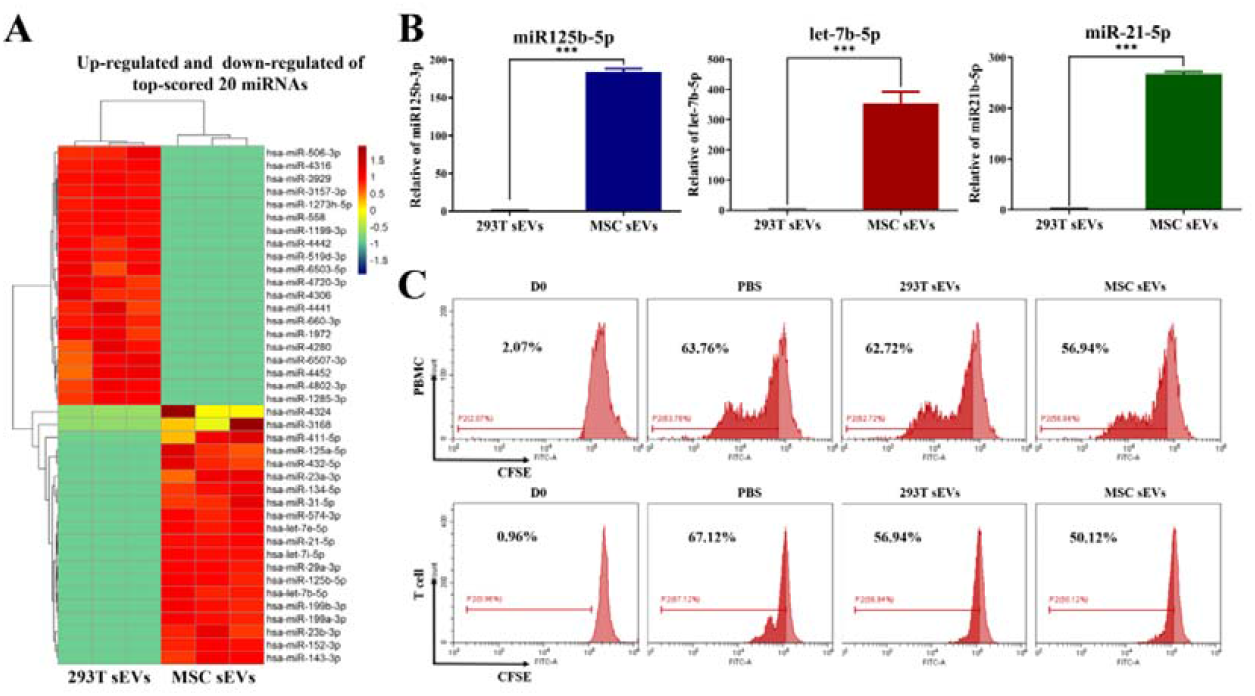
MSC sEVs has a negative immune regulation function. (related to Figure 2) A) Heat map of miRNAs targeting these immune genes created using 4,872 Immunologic Signature Gene Sets provided by the GSEA website, *n* = 3. B) Real time RT-PCR confirmation of miR125b-5p, let-7b-5p and miR-21-5p up-regulated by miRNA-sequencing, *n* = 3. Error bar, mean ± SEM. *P*-values are calculated using student T-test. \**P* < 0.05, \*\**P* < 0.01, \*\*\**P* < 0.001. C) Inhibition of PBMC and CD3^+^ T cells over a 5 day period by different sEVs groups using CFSE staining. PBMCs or T cells were stimulated with plate-bound CD3 and IL-2 (2 ng mL^-1^), then treated with the indicated different types of sEVs (50 μg mL^-1^). Cell proliferation was detected by CFSE at 5 days post-treatment, respectively. NC group: treated with PBS. D0 group: CFSE assay at 0 days post-treatment. CFSE staining was analyzed by flow cytometry.

**Figure S2.**
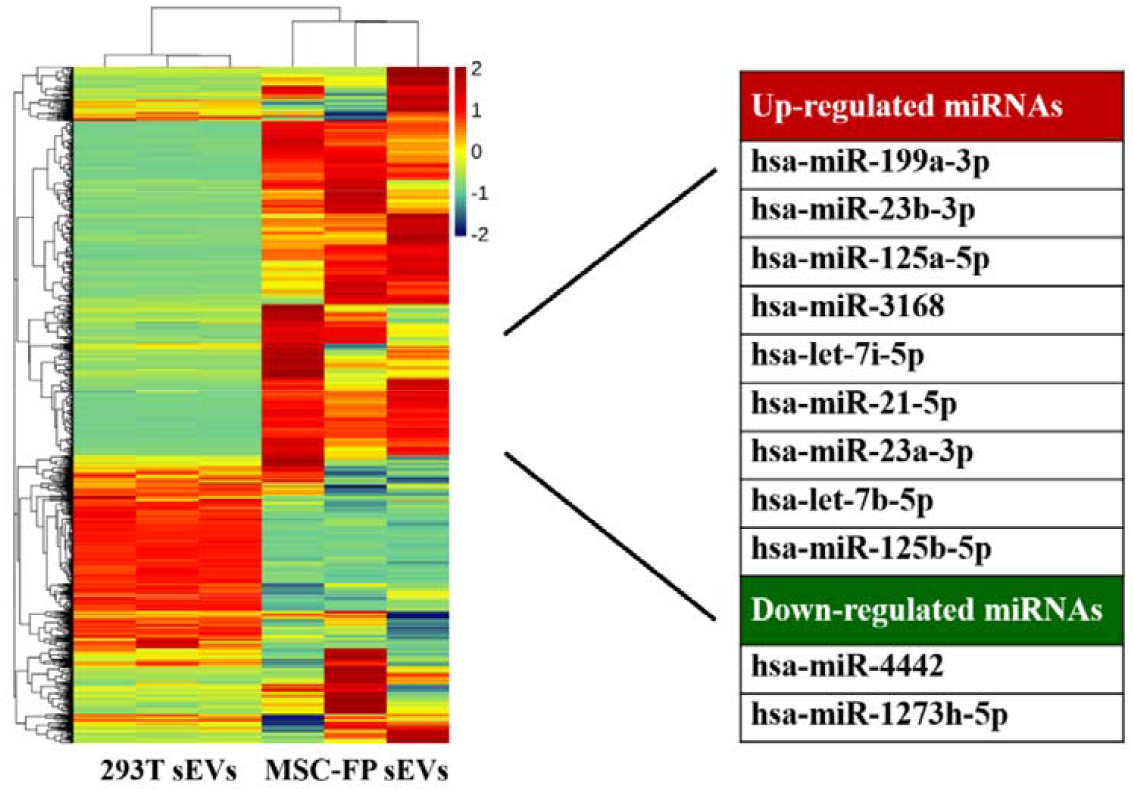
Modification couldn’t affect the negative immune regulation function of the miRNAs in exosomes. (related to Figure 2) Heat map of miRNAs in HEK-293T sEVs and MSC-FP sEVs targeting whole genome provided by the GSEA website, *n* = 3.

**Figure S3.**
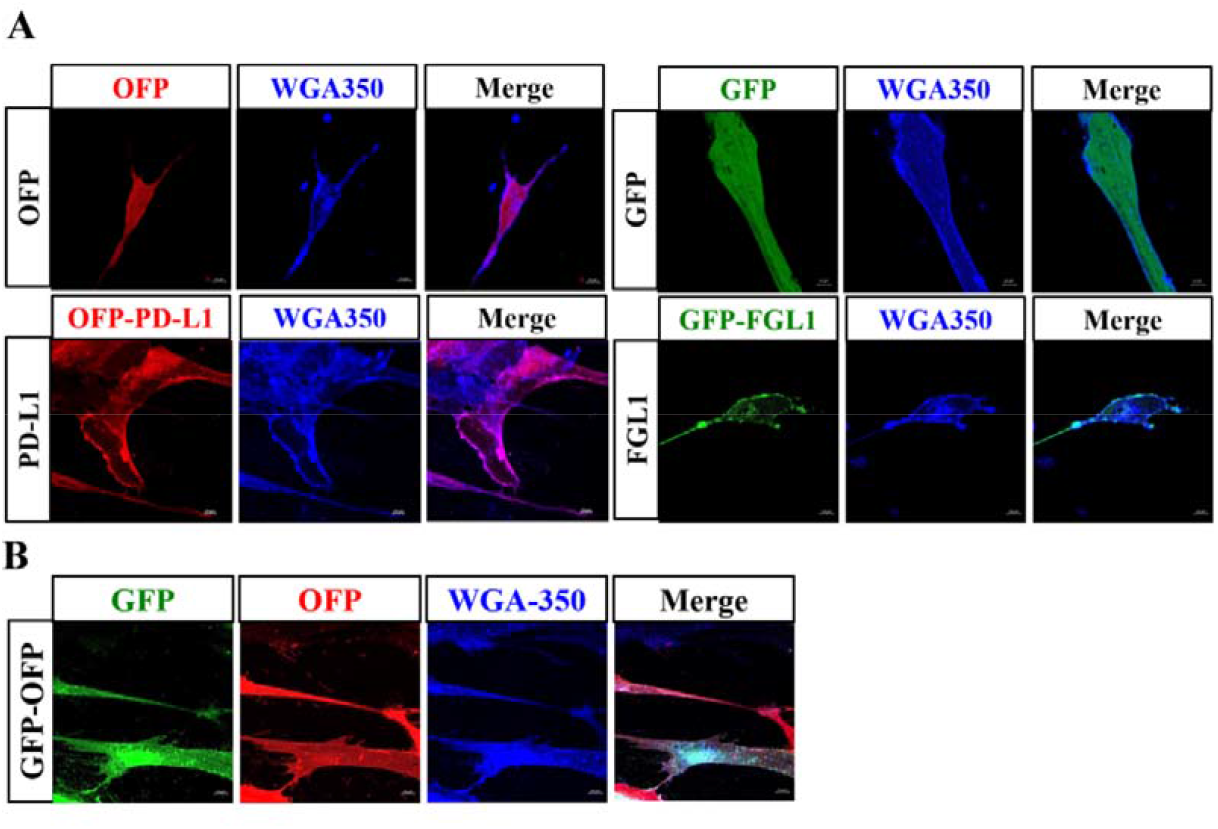
Construction of MSC cell line stably expressing GFP or OFP and GFP/OFP. (related to Figure 3) A) Confocal images indicated the expression of GFP or OFP in MSC cells. B) Confocal images indicated the expression of GFP and OFP in MSC cells. WGA-Alexa-350 was used to stain cell membranes. Scale bar: 10 *μ*m.

**Figure S4.**
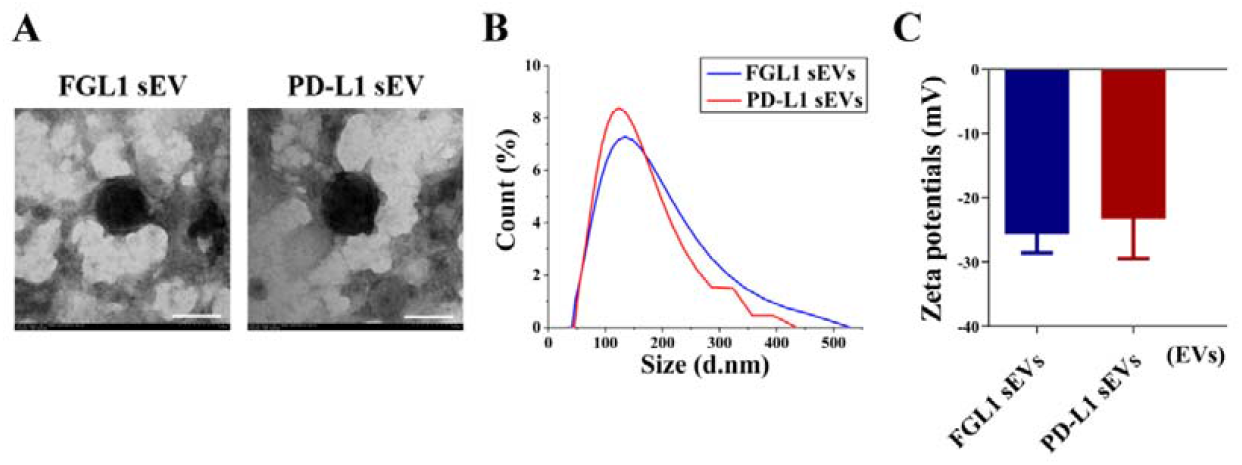
Establishment and characterization of FGL1 sEVs and PD-L1 sEVs. (related to Figure 3) A-C) The TEM images (A), size distribution (B), and the Zeta potential (C) of purified sEVs from MSC-FGL1 and MSC-PD-L1 cells, *n* = 3. (scale bar: 100 nm).

**Figure S5.**
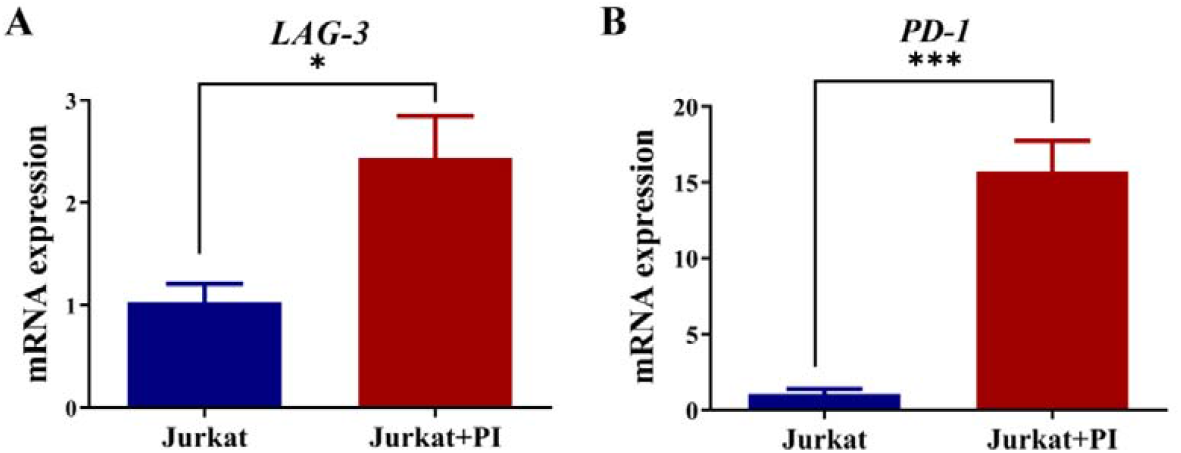
Upregulated PD-1 and LAG-3 mRNA levels of PBMC and Jurkat cell after PI stimulation. (related to Figure 4) A-B) Expression of LAG-3 (A) and PD-1 (B) mRNA levels in Jurkat cells upon P/I stimulation as detected by qPCR, *n* = 3. Error bar, mean ± SEM. *P*-values are calculated using student T-test. \**p*< 0.05, \*\**p*< 0.05, \*\*\**p*< 0.001.

**Figure S6.**
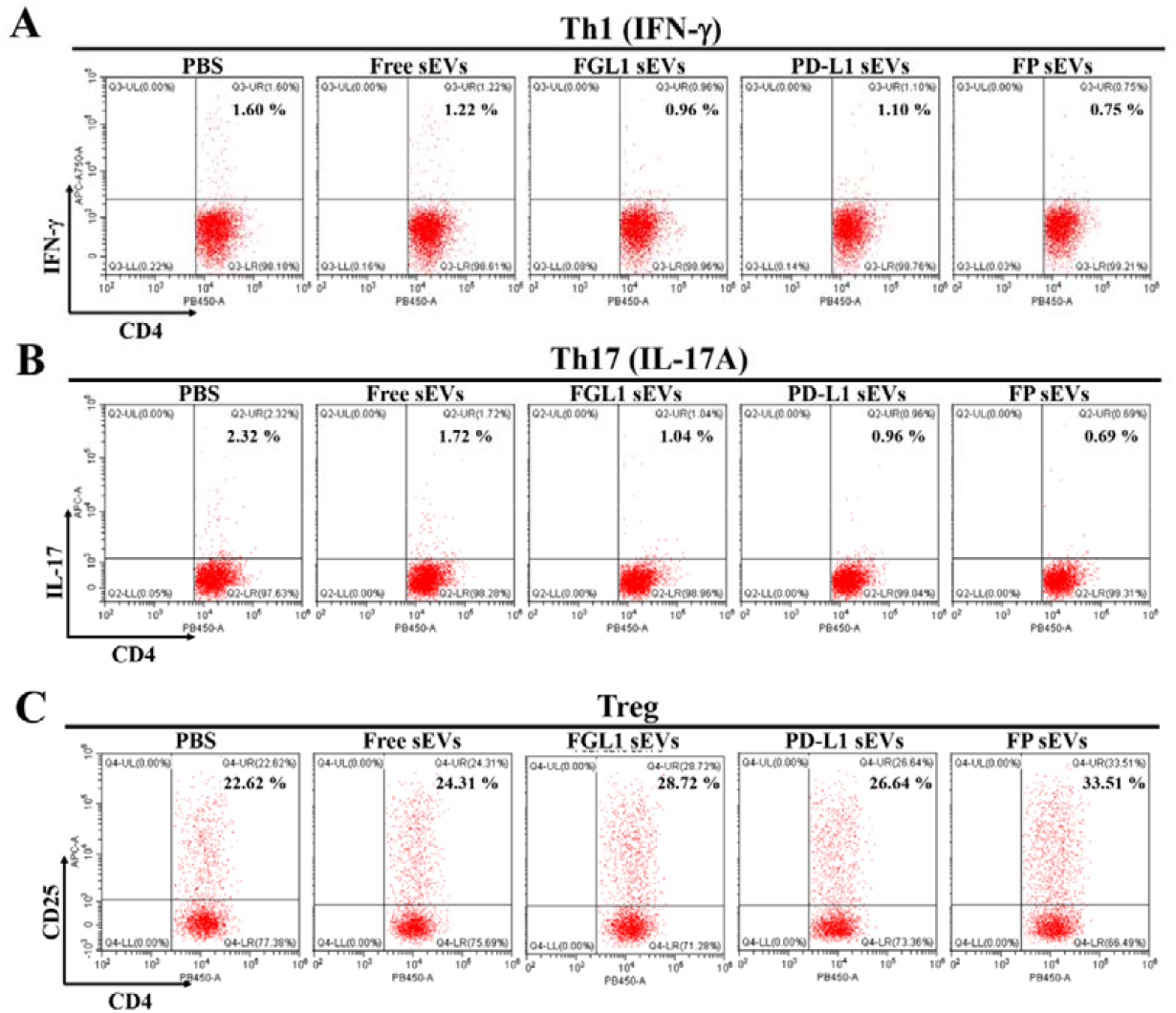
FGL1/PD-L1 sEVs could affect Th1, Th17 and Treg differentiation. (related to Figure 4) A-C) Flow cytometry analysis of CD4^+^ IFN-γ^+^ Th1 (A), CD4^+^ IL-17A^+^ Th17 (B) and CD4^+^ CD25^+^ Treg (C) expression in P/I-stimulated CD3^+^ T cells treated with or without different groups of sEVs.

**Figure S7.**
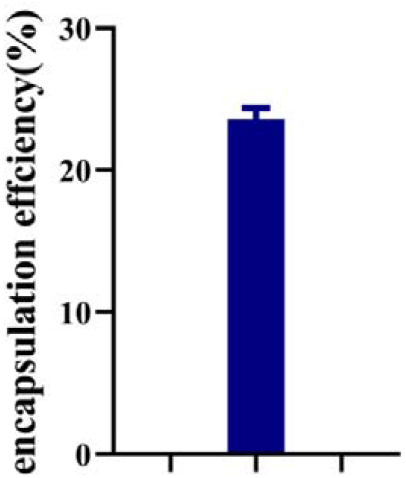
Encapsulation rate of FGL1/PD-L1 sEVs loaded with FK506. (related to Figure 4) FK506 was encapsulated in the FGL1/PD-L1 sEVs by electroporation, and its encapsulation rate was detected by UV-VIS at 300 nm (*n* = 3).

**Figure S8.**
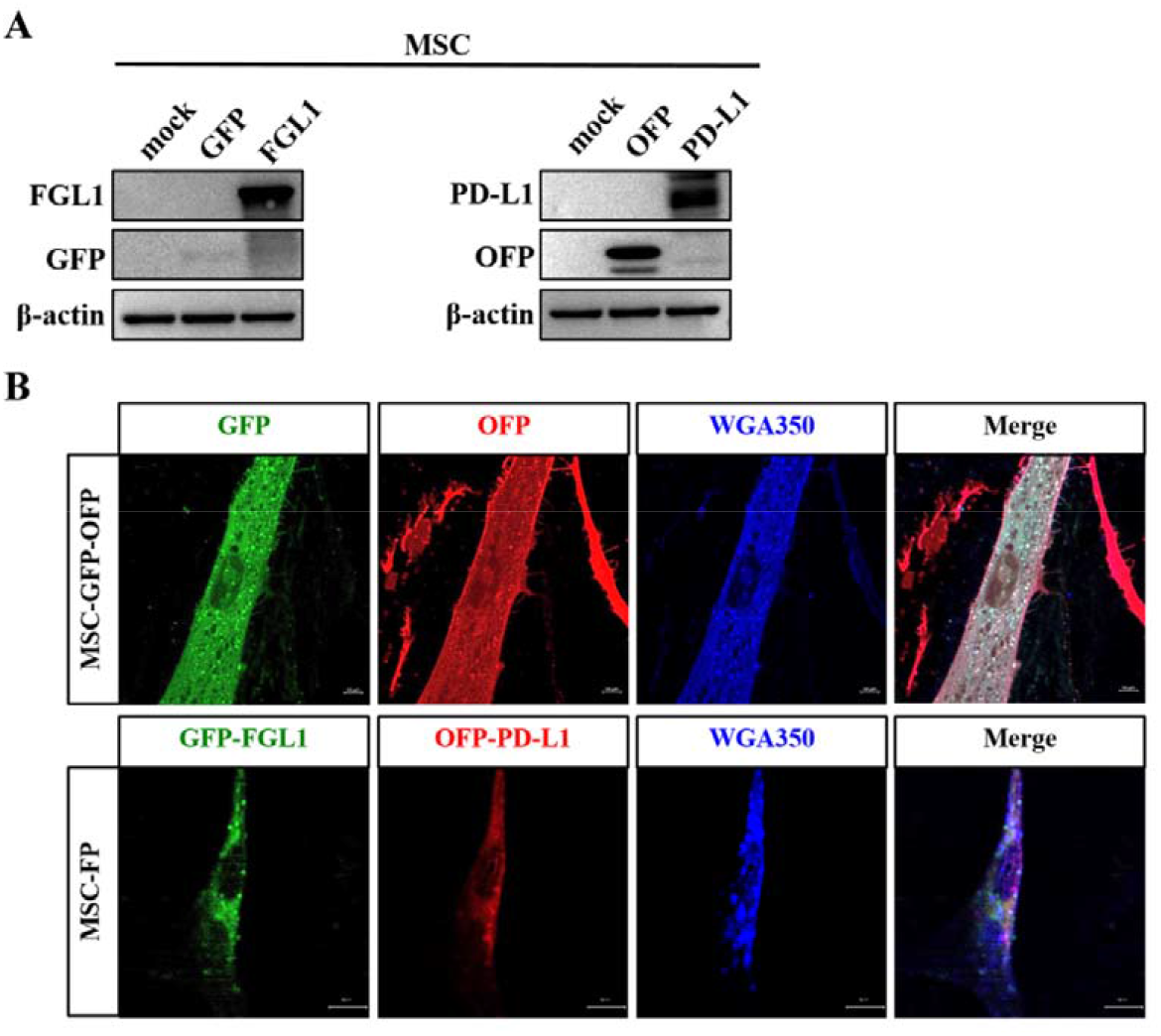
Establishment of MSC cells expressing mouse-FGL1/PD-L1. (related to Figure 5) A) Western blotting for mouse-PD-L1-OFP and mouse-FGL1-GFP in the whole cell lysate in MSCs. B) Confocal images indicate the expression of mouse-PD-L1-OFP, mouse-FGL1-GFP and mouse-PD-L1-OFP/mouse-FGL1-GFP in MSCs. Cell membranes stained with WGA Alexa 350. Scale bar: 10 *μ*m.

**Figure S9.**
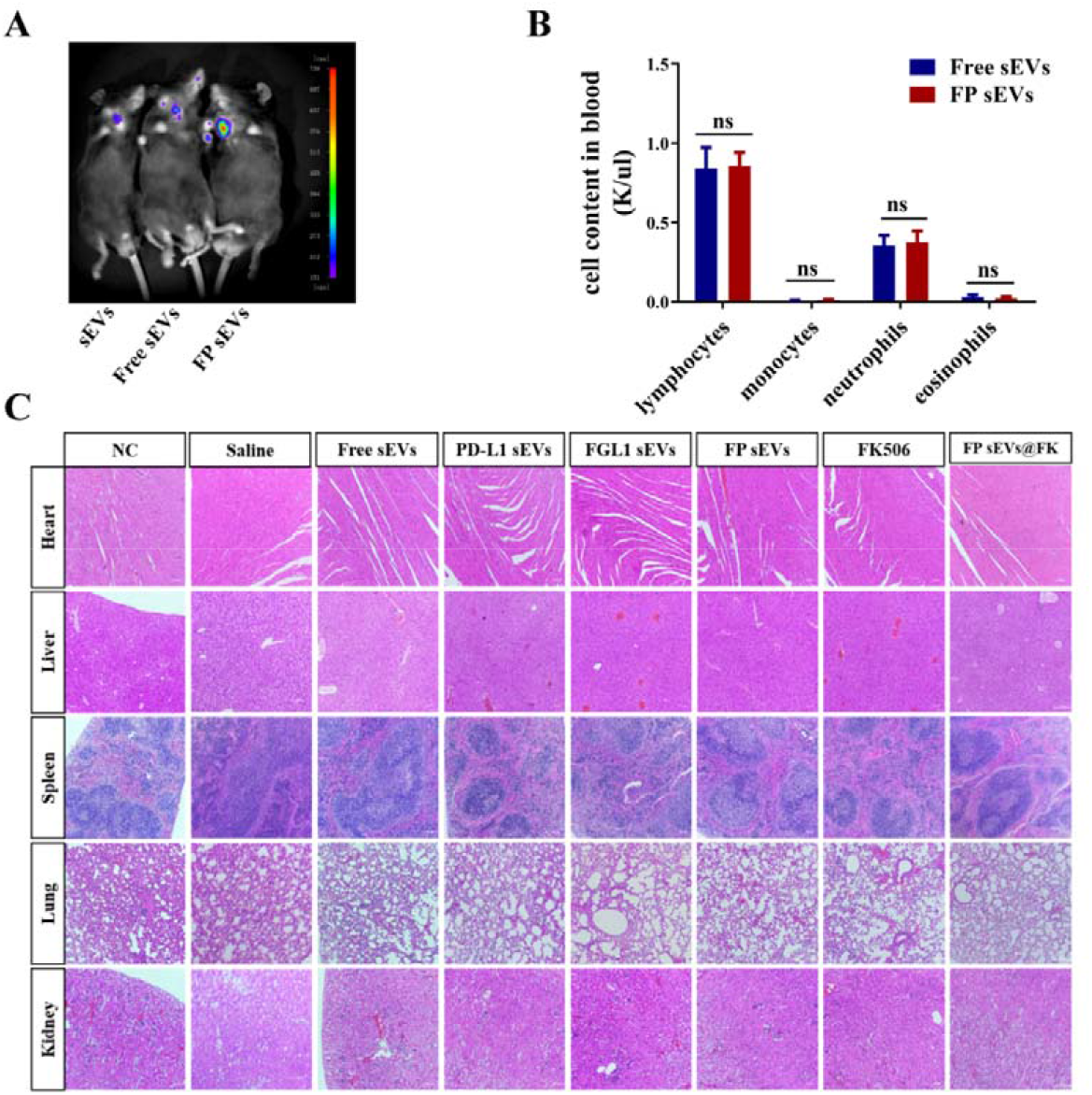
Bioluminescence imaging and toxicity test of FGL1/PD-L1 sEVs in mice. (related to Figure 5) A) *In vivo* Bioluminescence imaging of sEVs, Free sEVs, and FP sEVs originating from MSC cells via tail vein injection. sEVs: injected with MSC-sEVs. Free sEVs: injected with vector sEVs. B) Complete blood count test (CBC test). Mice were injected with Free sEVs or FP sEVs via tail vein injection for 14 days for whole blood analysis, *n* = 5. Error bar, mean ± SEM. *P*-values are calculated using student T-test. ns: not significant. C) Histological images obtained from the hearts, livers, spleens and kidneys of mice treated with NC, saline, or different groups of EVs for 14 days post-injection as compared to the NC group. Scale bar: 100 *μ*m.

**Figure S10.**
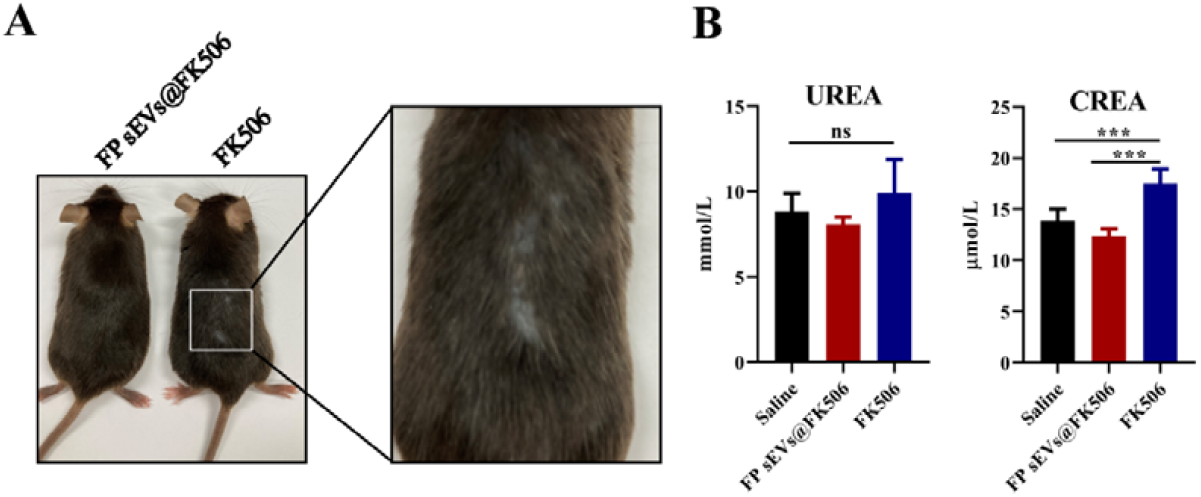
The appearance and biochemical analysis of FK506 and FP sEVs @ FK506 in heart allogenic mice. (related to Figure 5) A) Images highlighting the hair loss of heart allogenic mice after injecting with FK506 or FP sEVs@FK506. B) Blood samples were collected after injecting with saline, FK506 or FP sEVs@FK506 to measure creatinine (CREA) and urea nitrogen (UREA) by automatic blood analyzer. *n* = 5. Error bar, mean ± SEM. ns: not significant, \**P* < 0.05, \*\**P* < 0.01, \*\*\**P* < 0.001.

### Supplementary Tables

**Table S1.** Top-scored negative immunoregulation miRNAs between HEK-293T sEVs and MSC-FP sEVs.

| Up-regulated miRNAs | ID | P VALUE | Function |
| --- | --- | --- | --- |
| hsa-miR-199a-3p<sup>[27]</sup> | MIMAT0000232 | 1.45713E-08 | Inhibition of inflammation, immunity and resistance to kidney damage |
| hsa-miR-23b-3p<sup>[28]</sup> | MIMAT0000418 | 2.43588E-05 | Effective treatment of rheumatoid arthritis |
| hsa-miR-125a-5p<sup>[29]</sup> | MIMAT0000443 | 0.008456264 | Promote the generation of M2 type macrophages |
| hsa-miR-3168<sup>[30]</sup> | MIMAT0015043 | 8.46957E-25 | Suppress the immune system and relieve asthma |
| hsa-let-7i-5p<sup>[31]</sup> | MIMAT0000415 | 1.19388E-33 | Inhibition of IL - 6 |
| hsa-miR-21-5p<sup>[32]</sup> | MIMAT0000076 | 0.000657411 | Inhibits the immune function of dendritic cells |
| hsa-miR-23a-3p<sup>[33]</sup> | MIMAT0000078 | 3.35175E-05 | Upregulate the expression of PD-L1 to inhibit T cells function |
| hsa-let-7b-5p<sup>[34]</sup> | MIMAT0000063 | 4.70362E-27 | Inhibit the expression of inflammatory cytokines in macrophages |
| hsa-miR-125b-5p<sup>[35]</sup> | MIMAT0000423 | 1.37316E-88 | Inhibition of inflammation, immunity and resistance to kidney damage |
| Down-regulated miRNAs | ID | P VALUE | Function |
| hsa-miR-4442<sup>[36]</sup> | MIMAT0018960 | 1.2899E-122 | Downregulation inhibits inflammation and relieves dermatomyositis |
| hsa-miR-1273h-5p<sup>[37]</sup> | MIMAT0030415 | 1.0546E-166 | Increased in the PBMCs of SLE patients |

**Table S2.**
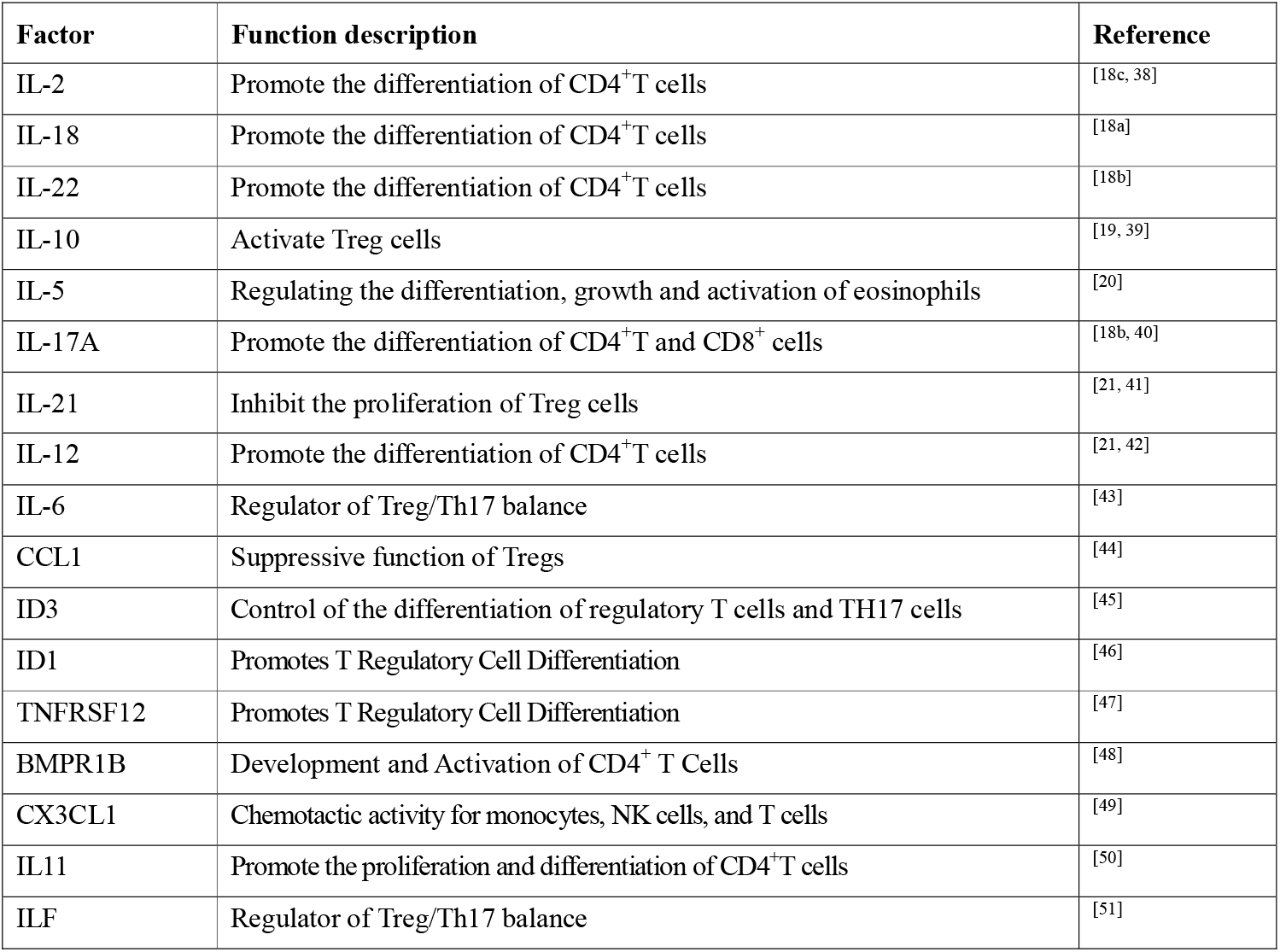
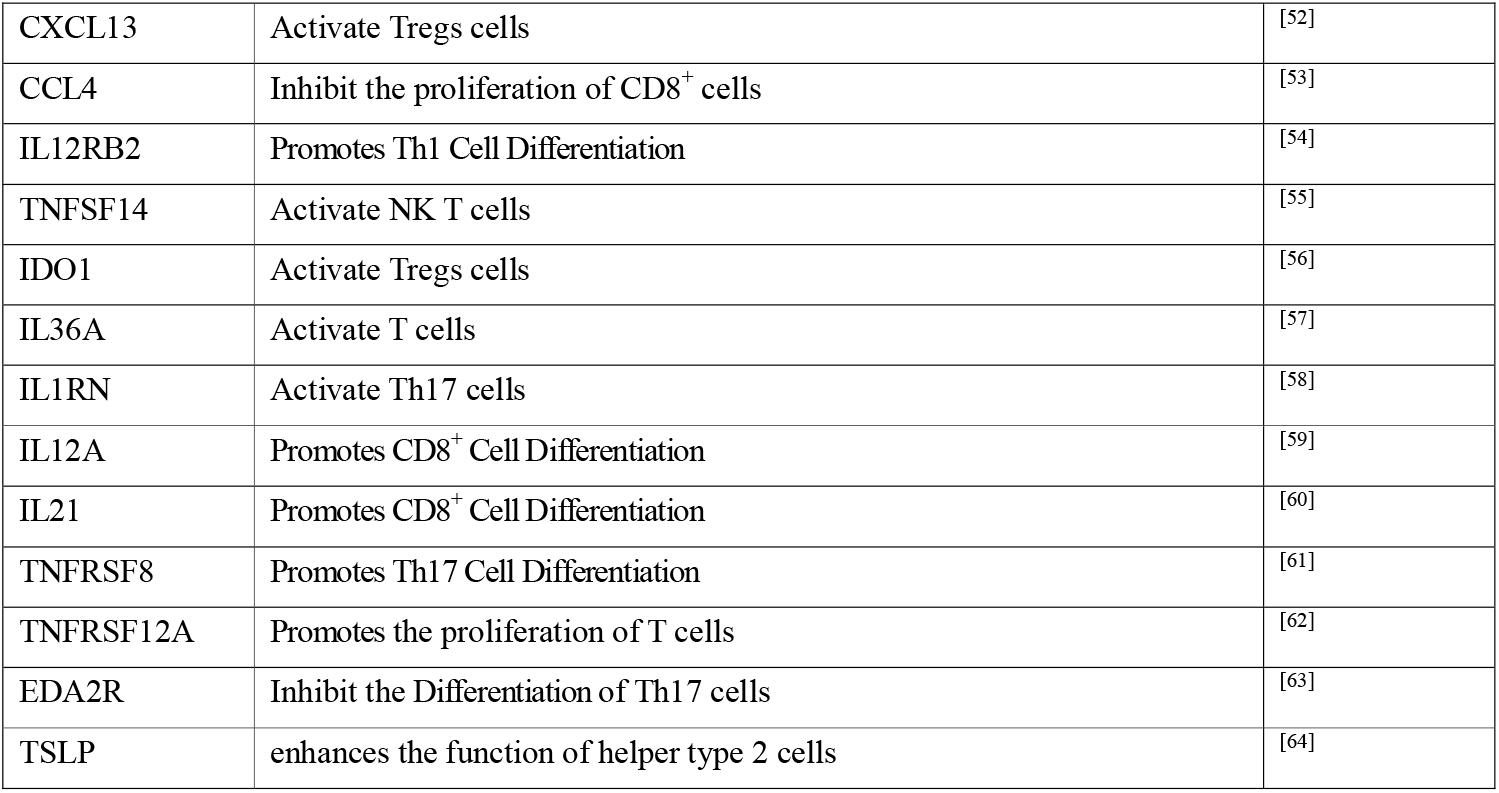
Description of T cell related factors.

**Table S3.**
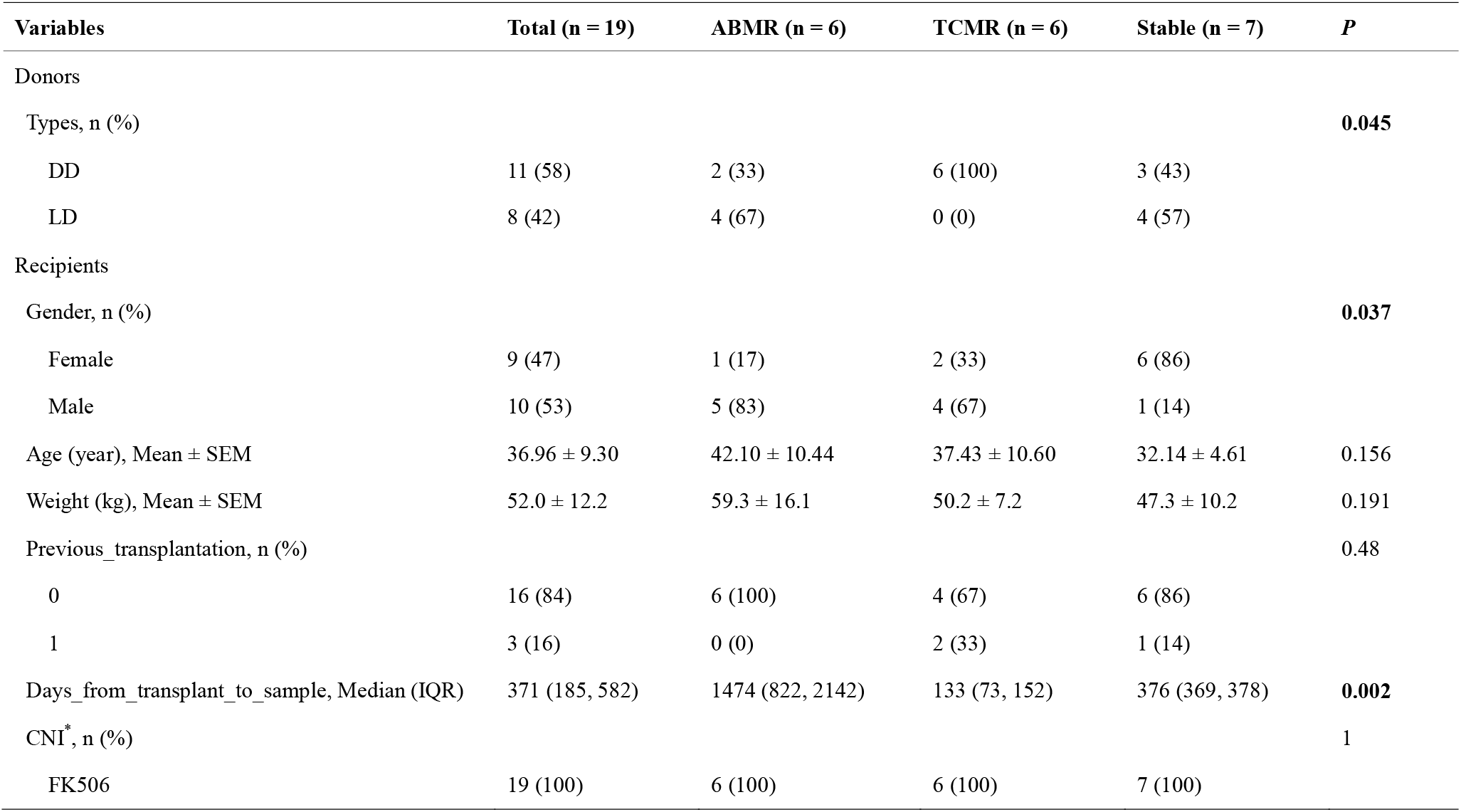
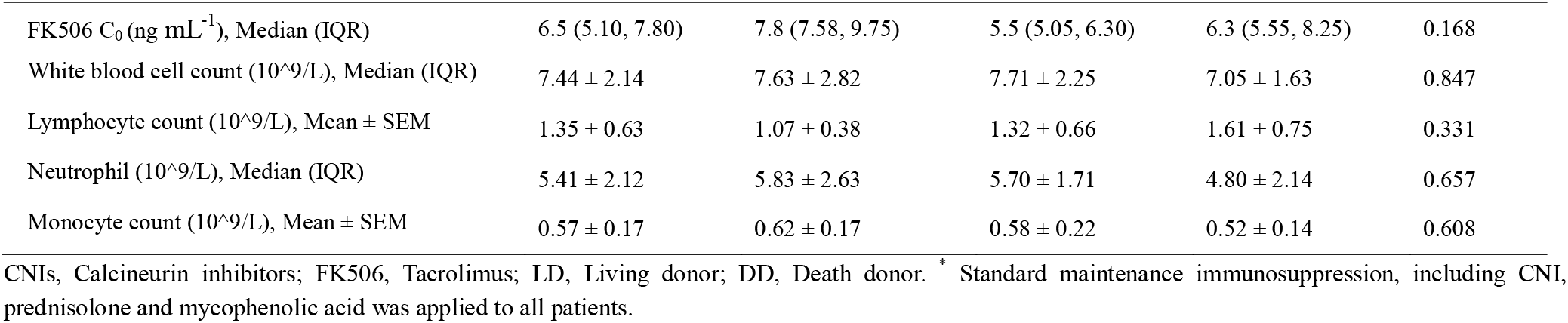
The demographic clinical characteristics in kidney transplant patients.

**Table S4.**
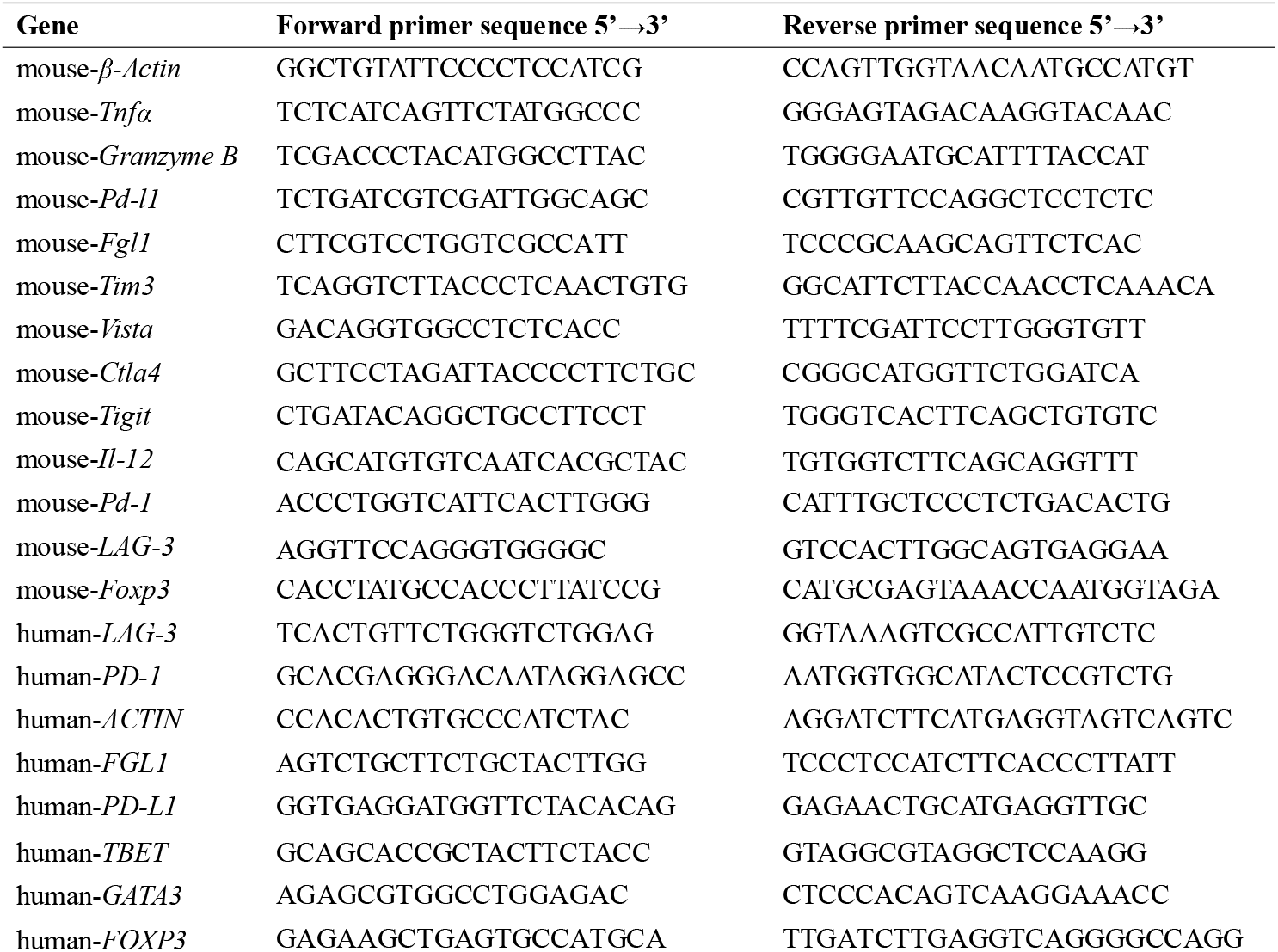
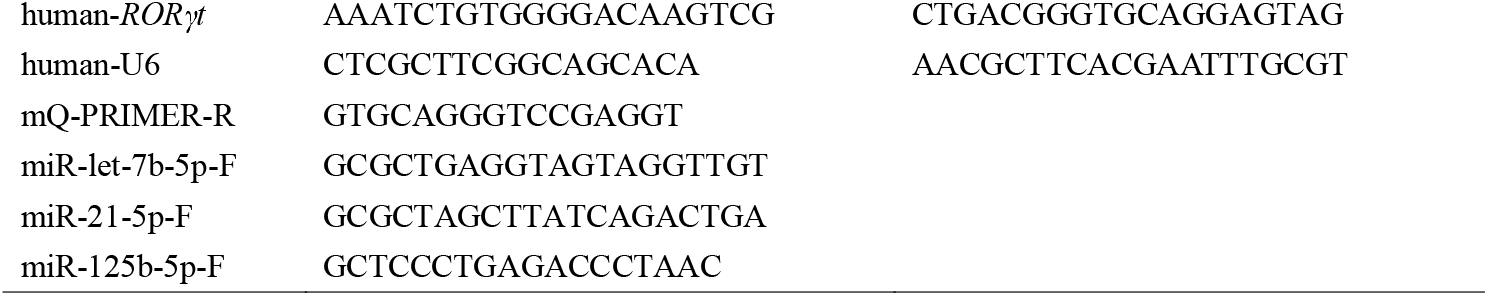
qPCR primer sequences.

## References

[1]a) F. Dumont, Curr. Med. Chem. 2000, 7 (7), 731;

b) F. Vincenti, S. C. Jensik, R. S. Filo, J. Miller, J. Pirsch, Transplantation 2002, 73 (5), 775;

c) U. S. M. F. L. S. Group, N. Engl. J. Med. 1994, 331 (17), 1110;

d) J.-U. Lee, L.-K. Kim, J.-M. Choi, Front. immunol. 2018, 9, 2747.

[2]a) R. I. Lechler, M. Sykes, A. W. Thomson, L. A. Turka, Nat. Med. 2005, 11 (6), 605;

b) M. D. Cahalan, Nat. Med. 2011, 17 (6), 662;

c) C. Y. Cheung, S. C. W. Tang, Nephrol. Dial. Transplant. 2019, 34 (6), 914;

d) R. Gioco, D. Corona, B. Ekser, L. Puzzo, G. Inserra, F. Pinto, C. Schipa, F. Privitera, P. Veroux, M. Veroux, World J. Gastroenterol. 2020, 26 (38), 5797.

[3]a) L. Long, X. Zhang, F. Chen, Q. Pan, P. Phiphatwatchara, Y. Zeng, H. Chen, cancer, Genes 2018, 9 (5-6), 176;

b) M. A. ElTanbouly, Y. Zhao, E. Nowak, J. Li, E. Schaafsma, I. Le Mercier, S. Ceeraz, J. L. Lines, C. Peng, C. Carriere, Science 2020, 367 (6475), 1;

c) L. Tu, R. Guan, H. Yang, Y. Zhou, W. Hong, L. Ma, G. Zhao, M. Yu, Int. J. Cancer. 2020, 147 (2), 423;

d) A. Assal, J. Kaner, G. Pendurti, X. Zang, Immunotherapy 2015, 7 (11), 1169.

[4] Z. Yin, M. Yu, T. Ma, C. Zhang, S. Huang, M. R. Karimzadeh, A. A. Momtazi-Borojeni, S. Chen, J. Immunother. Cancer. 2021, 9 (1), e001698.

[5]a) P. A. Ott, F. S. Hodi, C. Robert, Clin. Cancer. Res. 2013, 19 (19), 5300;

b) H. O. Alsaab, S. Sau, R. Alzhrani, K. Tatiparti, K. Bhise, S. K. Kashaw, A. K. Iyer, Front. Pharmacol. 2017, 8, 561;

c) Y. Jiang, X. Zhao, J. Fu, H. Wang, Front. Immunol. 2020, 11, 339.

[6] Y. Zhai, R. Moosavi, M. Chen, Front. Immunol. 2021, 12, 645699.

[7]a) H. Wang, Y. Liu, R. He, D. Xu, J. Zang, N. Weeranoppanant, H. Dong, Y. Li, Biomater. Sci. 2020, 8 (2), 552;

b) P. D. Robbins, A. Dorronsoro, C. N. Booker, J. Clin. Investig. 2016, 126 (4), 1173.

[8] Z. Xu, H.-i. Tsai, Y. Xiao, Y. Wu, D. Su, M. Yang, H. Zha, F. Yan, X. Liu, F. Cheng, ACS nano 2020, 14 (7), 7959.

[9]a) G. Abril-Rodriguez, A. Ribas, Cancer Cell 2017, 31 (6), 848;

b) J. B. Haanen, C. Robert, Prog. Tumor Res. 2015, 42, 55.

[10]a) X. Wang, H. Zhang, H. Yang, M. Bai, T. Ning, S. Li, J. Li, T. Deng, G. Ying, Y. Ba, Curr. Cancer Drug Tar. 2018, 18 (4), 347;

b) E. J. Bunggulawa, W. Wang, T. Yin, N. Wang, C. Durkan, Y. Wang, G. Wang, J. Nanotechnol. 2018, 16 (1), 1;

c) T. Yamamoto, N. Kosaka, T. Ochiya, Sci. Technol. Adv. Mat. 2019, 20 (1), 746;

d) J. Meldolesi, Curr. Biol. 2018, 28 (8), R435.

[11] G. Chen, A. C. Huang, W. Zhang, G. Zhang, M. Wu, W. Xu, Z. Yu, J. Yang, B. Wang, H. Sun, H. Xia, Q. Man, W. Zhong, L. F. Antelo, B. Wu, X. Xiong, X. Liu, L. Guan, T. Li, S. Liu, R. Yang, Y. Lu, L. Dong, S. McGettigan, R. Somasundaram, R. Radhakrishnan, G. Mills, Y. Lu, J. Kim, Y. H. Chen, H. Dong, Y. Zhao, G. C. Karakousis, T. C. Mitchell, L. M. Schuchter, M. Herlyn, E. J. Wherry, X. Xu, W. Guo, Nature 2018, 560 (7718), 382.

[12] A. G. Ramsay, Br. J. Haematol. 2013, 162 (3), 313.

[13]a) P. Lai, J. Weng, L. Guo, X. Chen, X. Du, Biomark. Res. 2019, 7 (1), 1;

b) H. K. Salem, C. Thiemermann, Stem Cells 2010, 28 (3), 585.

[14]a) P. Lai, X. Chen, L. Guo, Y. Wang, X. Liu, Y. Liu, T. Zhou, T. Huang, S. Geng, C. Luo, X. Huang, S. Wu, W. Ling, X. Du, C. He, J. Weng, J. Hematol. Oncol. 2018, 11 (1), 135;

b) L. Wang, Z. Gu, X. Zhao, N. Yang, F. Wang, A. Deng, S. Zhao, L. Luo, H. Wei, L. Guan, Z. Gao, Y. Li, L. Wang, D. Liu, C. Gao, Stem Cells Dev. 2016, 25 (24), 1874;

c) L. Kordelas, V. Rebmann, A. K. Ludwig, S. Radtke, J. Ruesing, T. R. Doeppner, M. Epple, P. A. Horn, D. W. Beelen, B. Giebel, Leukemia 2014, 28 (4), 970;

d) S. Farzamfar, A. Hasanpour, N. Nazeri, H. Razavi, M. Salehi, S. Shafei, V. T. Nooshabadi, A. Vaez, A. Ehterami, H. Sahrapeyma, J. Ai, J. Cell Physiol. 2019, 234 (8), 12290.

[15] D. Su, H.-I. Tsai, Z. Xu, F. Yan, Y. Wu, Y. Xiao, X. Liu, Y. Wu, S. Parvanian, W. Zhu, J. Extracell Vesicles 2020, 9 (1), 1709262.

[16] J. Wang, M. F. Sanmamed, I. Datar, T. T. Su, L. Ji, J. Sun, L. Chen, Y. Chen, G. Zhu, W. Yin, Cell 2019, 176 (1-2), 334.

[17]a) H. K. Lee, S. Finniss, S. Cazacu, C. Xiang, C. Brodie, Stem Cells Dev. 2014, 23 (23), 2851;

b) D. Ti, H. Hao, C. Tong, J. Liu, L. Dong, J. Zheng, Y. Zhao, H. Liu, X. Fu, W. Han, J. Transl. Med. 2015, 13 (1), 1.

[18]a) M. Sawada, T. Kawayama, H. Imaoka, Y. Sakazaki, H. Oda, S. Takenaka, Y. Kaku, K. Azuma, M. Tajiri, N. Edakuni, M. Okamoto, S. Kato, T. Hoshino, PLoS One 2013, 8 (1), e54623;

b) T. J. Scriba, B. Kalsdorf, D. A. Abrahams, F. Isaacs, J. Hofmeister, G. Black, H. Y. Hassan, R. J. Wilkinson, G. Walzl, S. J. Gelderbloem, H. Mahomed, G. D. Hussey, W. A. Hanekom, J. Immunol. 2008, 180 (3), 1962;

c) H. Abken, Transplantation 2021, 105 (7), 1394.

[19] J. Niu, W. Yue, Y. Song, Y. Zhang, X. Qi, Z. Wang, B. Liu, H. Shen, X. Hu, Clin. Exp. Immunol. 2014, 176 (3), 473.

[20] M. Goldman, A. Le Moine, M. Braun, V. Flamand, D. Abramowicz, Trends Immunol. 2001, 22 (5), 247.

[21] C. Jandl, S. M. Liu, P. F. Canete, J. Warren, W. E. Hughes, A. Vogelzang, K. Webster, M. E. Craig, G. Uzel, A. Dent, Nat. Commun. 2017, 8 (1), 1.

[22]a) Y. J. A. o. Bentata, Artif. Organs. 2020, 44 (2), 140;

b) H. i. Tsai, X. Zeng, L. Liu, S. Xin, Y. Wu, Z. Xu, H. Zhang, G. Liu, Z. Bi, D. Su, EMBO Mol. Med. 2021, 13 (3), e12834.

[23] R. Costello, A. Kissenpfennig, P. N. Martins, J. McDaid, Expert Opin. Drug Discov. 2018, 13 (11), 1041.

[24] R.-Y. Huang, C. Eppolito, S. Lele, P. Shrikant, J. Matsuzaki, K. Odunsi, Oncotarget 2015, 6 (29), 27359.

[25]a) M. Żmigrodzka, M. Guzera, A. Miśkiewicz, D. Jagielski, A. Winnicka, Tumor Biol. 2016, 37 (11), 14391;

b) X. Zhang, C. Wang, J. Wang, Q. Hu, B. Langworthy, Y. Ye, W. Sun, J. Lin, T. Wang, J. Fine, Adv. Mater. 2018, 30 (22), 1707112;

c) X. Zhang, J. Wang, Z. Chen, Q. Hu, C. Wang, J. Yan, G. Dotti, P. Huang, Z. Gu, Nano Lett. 2018, 18 (9), 5716;

d) S. Tan, T. Wu, D. Zhang, Z. Zhang, Theranostics 2015, 5 (8), 863;

e) S. Ghosh, K. Girigoswami, A. Girigoswami, Nanomedicine (Lond) 2019, 14 (15), 2067.

[26]a) G. Xu, L. Wang, W. Chen, F. Xue, X. Bai, L. Liang, X. Shen, M. Zhang, D. Xia, T. Liang, Liver Transpl. 2010, 16 (3), 357;

b) A. Y. Rudensky, M. Gavin, Y. Zheng, Cell 2006, 126 (2), 253.

[27] G. Zhu, L. Pei, F. Lin, H. Yin, X. Li, W. He, N. Liu, X. Gou, J. Cell Physiol. 2019, 234(12), 23736.

[28] R. Li, Q. Ruan, F. Yin, K. Zhao, J. Pharmacol. Sci. 2021, 145 (1), 69.

[29] S. Banerjee, H. Cui, N. Xie, Z. Tan, S. Yang, M. Icyuz, V. J. Thannickal, E. Abraham, G. Liu, J. Biol. Chem. 2013, 288 (49), 35428.

[30] T. Bahmer, S. Krauss-Etschmann, D. Buschmann, J. Behrends, H. Watz, A. M. Kirsten, F. Pedersen, B. Waschki, O. Fuchs, M. W. Pfaffl, E. von Mutius, K. F. Rabe, G. Hansen, M. V. Kopp, I. R. Konig, S. Bartel, Allergy 2021, 76 (1), 366.

[31] X. Wang, H. X. Wang, Y. L. Li, C. C. Zhang, C. Y. Zhou, L. Wang, Y. L. Xia, J. Du, H. H. Li, Hypertension 2015, 66 (4), 776.

[32] M. Reis, E. Mavin, L. Nicholson, K. Green, A. M. Dickinson, X. N. Wang, Front. Immunol. 2018, 9, 2538.

[33] J. Liu, L. Fan, H. Yu, J. Zhang, Y. He, D. Feng, F. Wang, X. Li, Q. Liu, Y. Li, Z. Guo, B. Gao, W. Wei, H. Wang, G. Sun, Hepatology 2019, 70 (1), 241.

[34] S. E. Nematian, R. Mamillapalli, T. S. Kadakia, M. Majidi Zolbin, S. Moustafa, H. S. Taylor, J. Clin. Endocrinol. Metab. 2018, 103 (1), 64.

[35] J. Y. Cao, B. Wang, T. T. Tang, Y. Wen, Z. L. Li, S. T. Feng, M. Wu, D. Liu, D. Yin, K. L. Ma, R. N. Tang, Q. L. Wu, H. Y. Lan, L. L. Lv, B. C. Liu, Theranostics 2021, 11 (11), 5248.

[36] Y. Y. Yang, X. X. Zuo, H. L. Zhu, S. J. Liu, Beijing Da Xue Xue Bao Yi Xue Ban 2019, 51 (2), 374.

[37] G. Guo, H. Wang, X. Shi, L. Ye, K. Wu, K. Lin, S. Ye, B. Li, H. Zhang, Q. Lin, S. Ye, X. Xue, C. Chen, J. Transl. Med. 2018, 16 (1), 370.

[38] S. Sad, T. R. Mosmann, J. Immunol. 1994, 153 (8), 3514.

[39] C. I. Kingsley, M. Karim, A. R. Bushell, K. J. Wood, J. Immunol. 2002, 168 (3), 1080.

[40] M. P. Crawford, S. Sinha, P. S. Renavikar, N. Borcherding, N. J. Karandikar, Proc. Natl. Acad. Sci. U. S. A. 2020, 117 (32), 19408.

[41] A. Suto, D. Kashiwakuma, S. Kagami, K. Hirose, N. Watanabe, K. Yokote, Y. Saito, T. Nakayama, M. J. Grusby, I. Iwamoto, H. Nakajima, J. Exp. Med. 2008, 205 (6), 1369.

[42] G. Trinchieri, S. Pflanz, R. A. Kastelein, Immunity 2003, 19 (5), 641.

[43] A. Kimura, T. Kishimoto, Eur. J. Immunol. 2010, 40 (7), 1830.

[44] D. B. Hoelzinger, S. E. Smith, N. Mirza, A. L. Dominguez, S. Z. Manrique, J. Lustgarten, J. Immunol. 2010, 184 (12), 6833.

[45] T. Maruyama, J. Li, J. P. Vaque, J. E. Konkel, W. Wang, B. Zhang, P. Zhang, B. F. Zamarron, D. Yu, Y. Wu, Y. Zhuang, J. S. Gutkind, W. Chen, Nat. Immunol. 2011, 12 (1), 86.

[46] C. Liu, H. C. Wang, S. Yu, R. Jin, H. Tang, Y. F. Liu, Q. Ge, X. H. Sun, Y. Zhang, J. Immunol. 2014, 193 (2), 663.

[47] H. Nishikii, B. S. Kim, Y. Yokoyama, Y. Chen, J. Baker, A. Pierini, M. Alvarez, M. Mavers, K. Maas-Bauer, Y. Pan, S. Chiba, R. S. Negrin, Blood 2016, 128 (24), 2846.

[48] M. Kuczma, P. Kraj, Vitam. Horm. 2015, 99, 171.

[49] B. A. Jones, M. Beamer, S. Ahmed, Mol. Interv. 2010, 10 (5), 263.

[50] A. Curti, A. Tafuri, M. R. Ricciardi, P. Tazzari, M. T. Petrucci, M. Fogli, M. Ratta, R. Lapalombella, E. Ferri, S. Tura, M. Baccarani, R. M. Lemoli, Haematologica 2002, 87 (4), 373.

[51] S. M. Metcalfe, Genes Immun. 2011, 12 (3), 157.

[52] B. P. Lee, W. Chen, H. Shi, S. D. Der, R. Forster, L. Zhang, J. Immunol. 2006, 176 (9), 5276.

[53] S. A. Joosten, K. E. van Meijgaarden, N. D. Savage, T. de Boer, F. Triebel, A. van der Wal, E. de Heer, M. R. Klein, A. Geluk, T. H. Ottenhoff, Proc. Natl. Acad. Sci. U. S. A. 2007, 104 (19), 8029.

[54] Z. Y. Zhou, S. L. Chen, N. Shen, Y. Lu, Autoimmun Rev. 2012, 11 (10), 699.

[55] F. Shi, Y. Xiong, Y. Zhang, C. Qiu, M. Li, A. Shan, Y. Yang, B. Li, Inflammation 2018, 41 (3), 1021.

[56] A. de Luca, S. Bozza, T. Zelante, S. Zagarella, C. D’Angelo, K. Perruccio, C. Vacca, A. Carvalho, C. Cunha, F. Aversa, L. Romani, Cell Mol. Immunol. 2010, 7 (6), 459.

[57] S. Abdollahi-Roodsaz, L. A. Joosten, M. I. Koenders, I. Devesa, M. F. Roelofs, T. R. Radstake, M. Heuvelmans-Jacobs, S. Akira, M. J. Nicklin, F. Ribeiro-Dias, W. B. van den Berg, J. Clin. Invest. 2008, 118 (1), 205.

[58] J. C. Waite, D. Skokos, Int. J. Inflam. 2012, 2012, 819467.

[59] N. Takemoto, A. M. Intlekofer, J. T. Northrup, E. J. Wherry, S. L. Reiner, J. Immunol. 2006, 177 (11), 7515.

[60] C. S. Hinrichs, R. Spolski, C. M. Paulos, L. Gattinoni, K. W. Kerstann, D. C. Palmer, C. A. Klebanoff, S. A. Rosenberg, W. J. Leonard, N. P. Restifo, Blood 2008, 111 (11), 5326.

[61] X. Sun, H. Yamada, K. Shibata, H. Muta, K. Tani, E. R. Podack, Y. Yoshikai, J. Immunol. 2010, 185 (4), 2222.

[62] L. Xia, L. Jiang, Y. Chen, G. Zhang, L. Chen, Cytokine 2021, 155658.

[63] P. G. Miller, M. B. Bonn, S. C. McKarns, J. Immunol. 2015, 195 (6), 2633.

[64] M. Kitajima, H. C. Lee, T. Nakayama, S. F. Ziegler, Eur. J. Immunol. 2011, 41 (7), 1862.

