## Supplemental file for "Engineered small extracellular vesicles as a FGL1/PD-L1 dual-targeting delivery system for alleviating immune rejection"

Dr. H.I. Tsai^[+]^, Y. Y. Wu, X. Y. Liu, Z. X. Xu, L. L. Wang, W. X. Zhang, D. D. Su, Prof. F. Cheng, Prof. H. B. Chen

School of Pharmaceutical Sciences (Shenzhen)

Sun Yat-sen University

Shenzhen 518107, P.R. China

Prof. H.T. Zhu, Prof. D.Q. Wang

Department of Medical Imaging

The Affiliated Hospital of Jiangsu University

Zhenjiang 212001, China

Dr. H. X. Zhang, Prof. L. S. Liu, Prof. C. X. Wang

Organ Transplant Center

The First Affiliated Hospital, Sun Yat-sen University

58 Zhongshan 2nd Road, Guangzhou, Guangdong 510080, China

Dr. Y. S. Huang, Dr. B. Jia

Department of Oral Surgery

Stomatological Hospital, Southern Medical University

Guangzhou 510280, Guangdong, PR China

Dr. F. U. Khan

Zoology Department UST

Bannu Kp 28100, Pakistan

Dr. X. F. Zhu, Prof. Y.X Pang

School of Traditional Medicine Materials Resource, Guangdong Pharmaceutical University Yunfu 527322, Guangdong, China.

Prof. R.Y. Yang

Department of Dermatology, The Seventh Medical Center of PLA General Hospital,

Peking 100010, China

Prof. J. Eriksson

Cell Biology, Biosciences, Faculty of Science and Engineering

Åbo Akademi University,

FI-20520, Turku, Finland

^[+]^ Present address: Department of Medical Imaging

The Affiliated Hospital of Jiangsu University

Zhenjiang 212001, China

H. Tsai, Y. Y. Wu and X. Y. Liu contributed equally to this work.

B. Jia, F. Cheng, and H. B. Chen are corresponding authors

**Supporting Information**

**Supplement Figures**

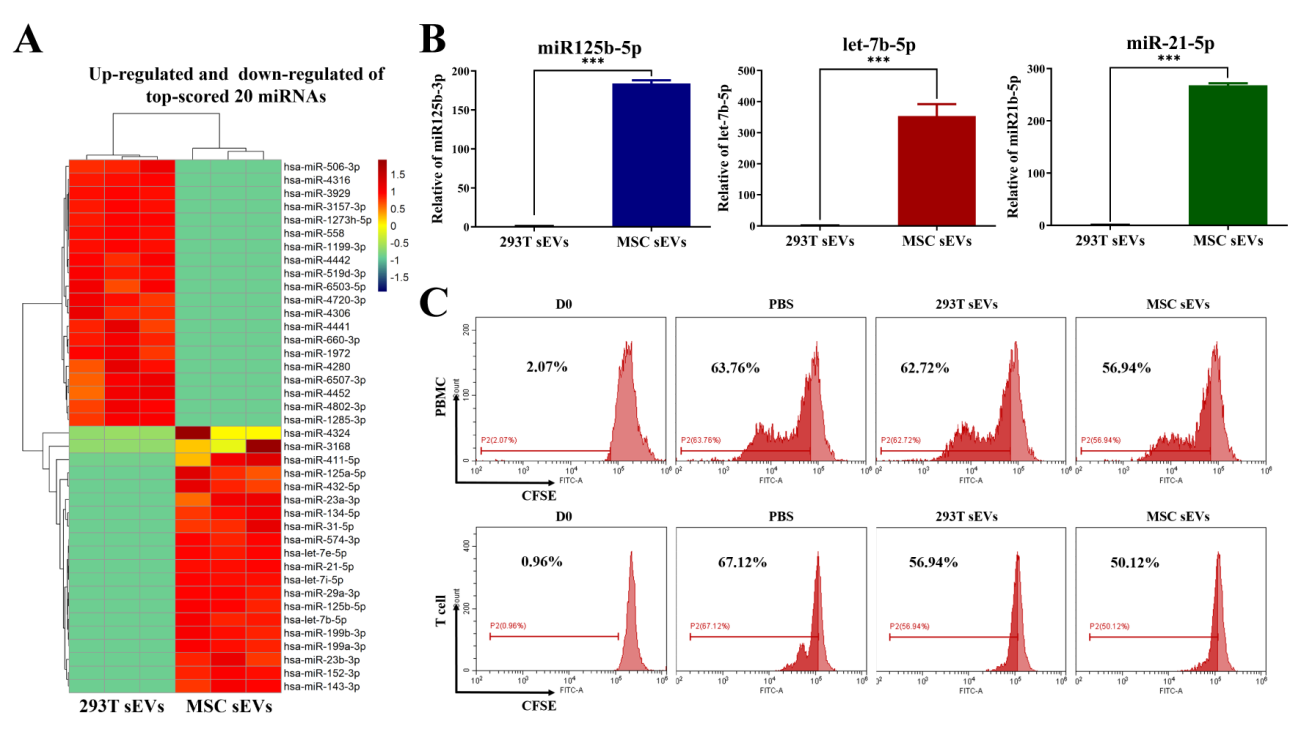

**Figure S1. MSC sEVs has a negative immune regulation function**. (related to Figure 2)

A) Heat map of miRNAs targeting these immune genes created using 4,872 Immunologic Signature Gene Sets provided by the GSEA website, *n* = 3. B) Real time RT-PCR confirmation of miR125b-5p, let-7b-5p and miR-21-5p up-regulated by miRNA-sequencing, *n* = 3. Error bar, mean ± SEM. *P*-values are calculated using student T-test. **P* < 0.05, ***P* < 0.01, ****P* < 0.001**.** C) Inhibition of PBMC and CD3^+^ T cells over a 5 day period by different sEVs groups using CFSE staining. PBMCs or T cells were stimulated with plate-bound CD3 (10 μg/mL) and IL-2 (2 ng/mL), then treated with the indicated different types of sEVs (50 μg/mL). Cell proliferation was detected by CFSE at 5 days post-treatment, respectively. NC group: treated with PBS. D0 group: CFSE assay at 0 days post-treatment. CFSE staining was analyzed by flow cytometry.

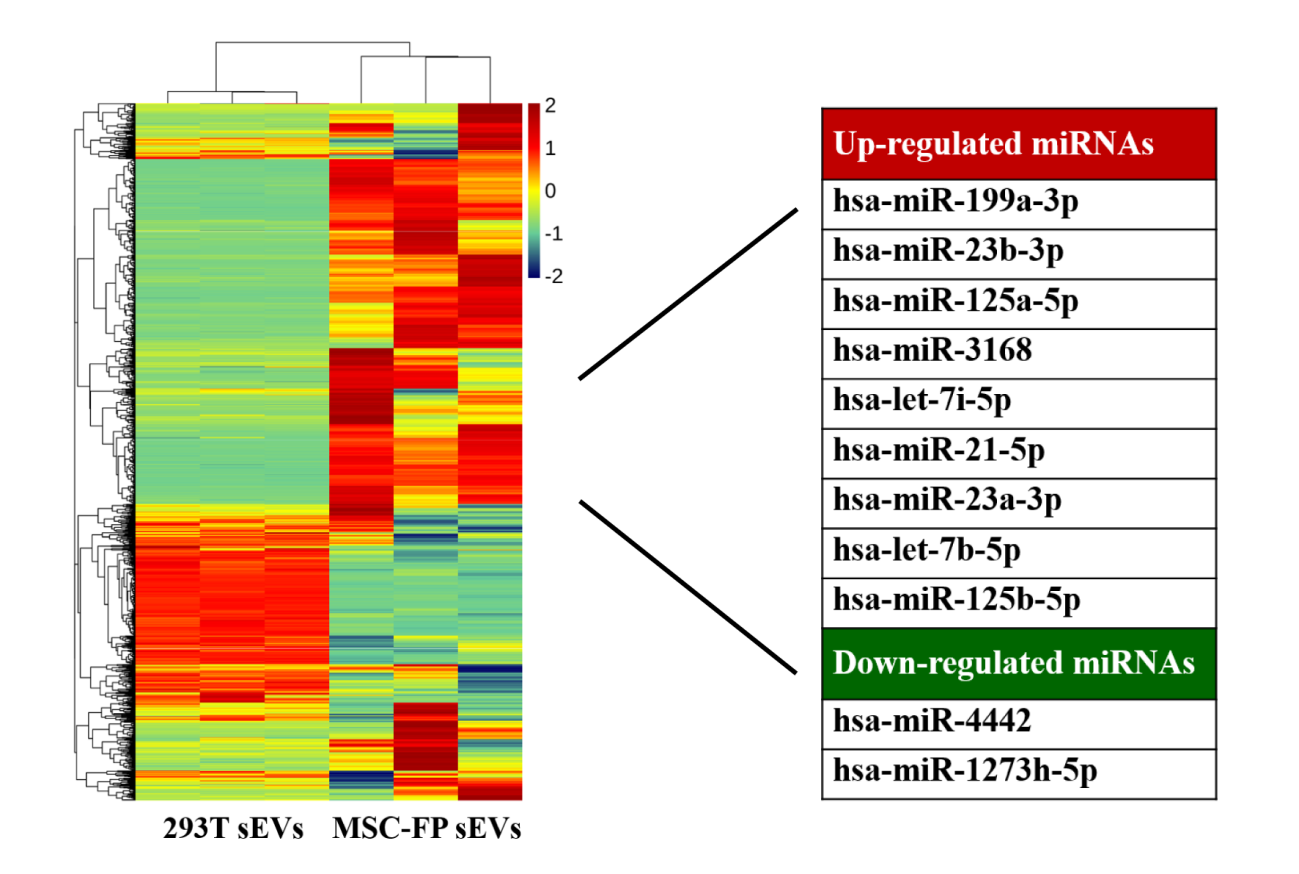

**Figure S2. Modification couldn’t affect the negative immune regulation function of the miRNAs in exosomes.** (related to Figure 2)

Heat map of miRNAs in HEK-293T sEVs and MSC-FP sEVs targeting whole genome provided by the GSEA website, *n* = 3.

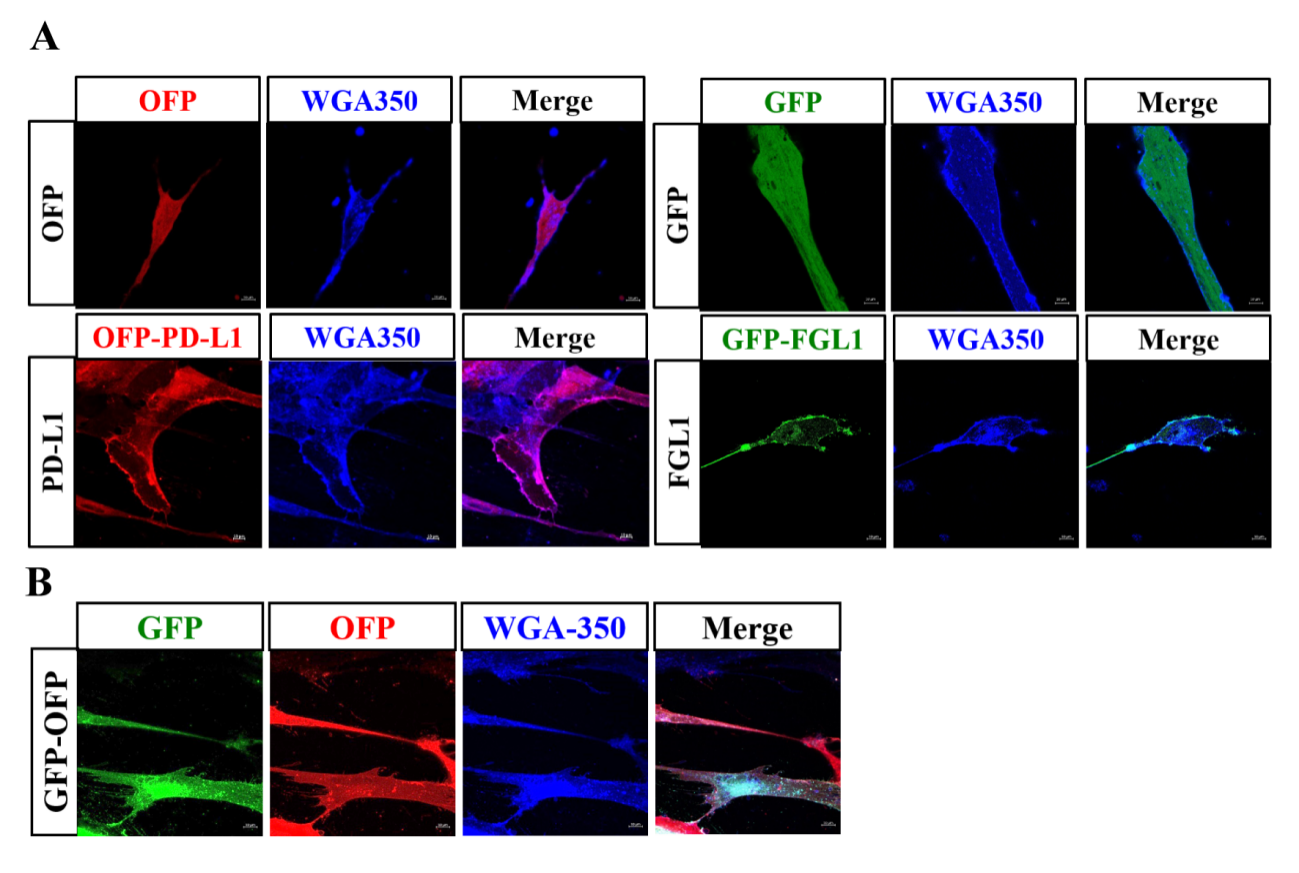

**Figure S3. Construction of MSC cell line stably expressing GFP or OFP and GFP/OFP**. (related to Figure 3)

A) Confocal images indicated the expression of GFP or OFP in MSC cells. B) Confocal images indicated the expression of GFP and OFP in MSC cells. WGA-Alexa-350 was used to stain cell membranes. Scale bar: 10 *µ*m.

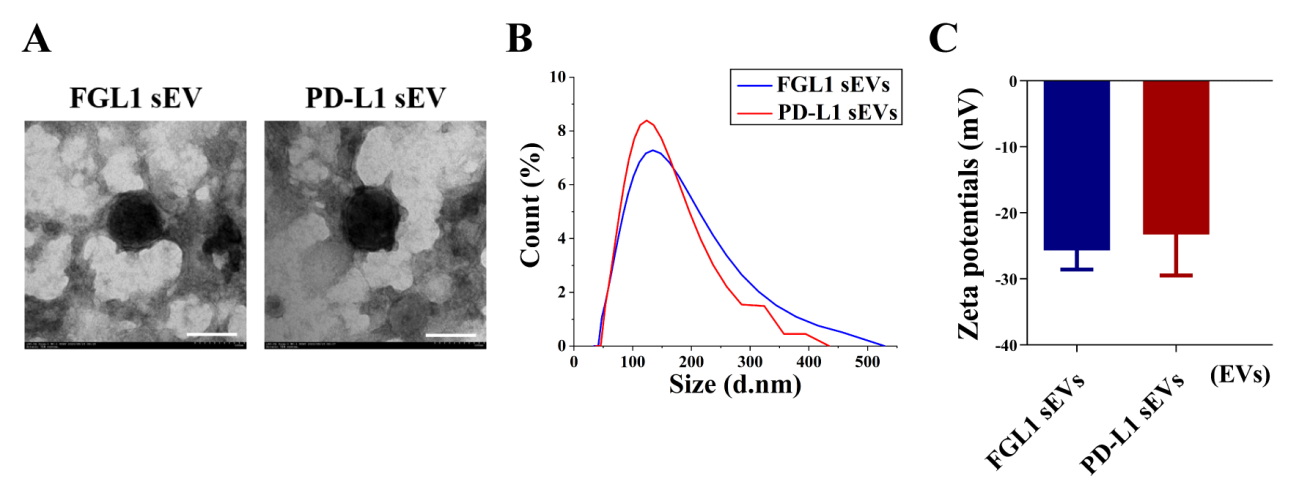

**Figure S4. Establishment and characterization of FGL1 sEVs and PD-L1 sEVs.** (related to Figure 3)

A-C) The TEM images (A), size distribution (B), and the Zeta potential (C) of purified sEVs from MSC-FGL1 and MSC-PD-L1 cells, *n* = 3. (scale bar: 100 nm).

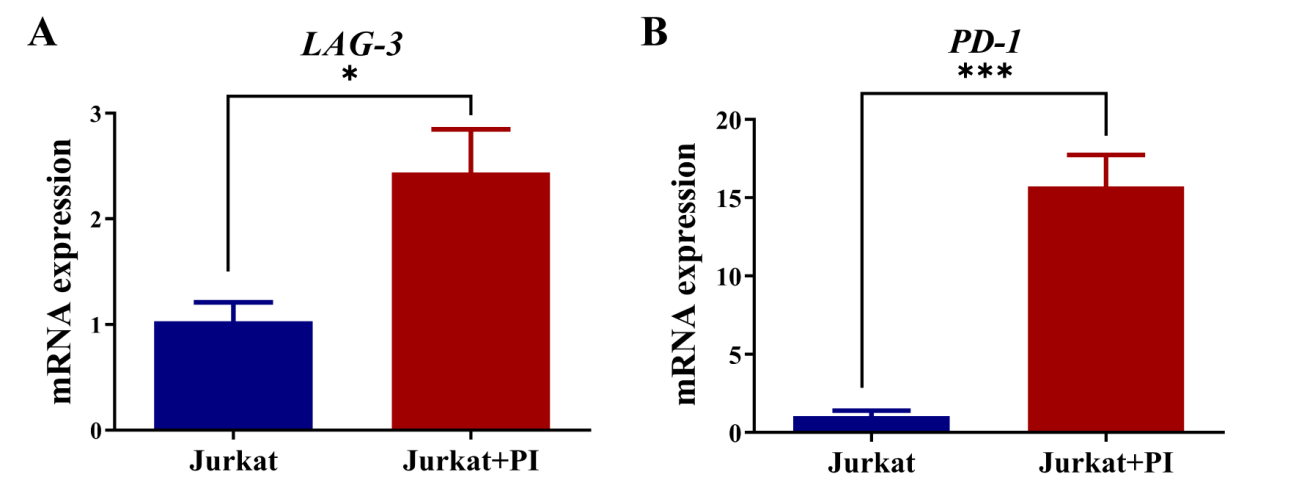

**Figure S5. Upregulated PD-1 and LAG-3 mRNA levels of PBMC and Jurkat cell after PI stimulation.** (related to Figure 4)

A-B) Expression of LAG-3 (A) and PD-1 (B) mRNA levels in Jurkat cells upon P/I stimulation as detected by qPCR, *n* = 3. Error bar, mean ± SEM. *P*-values are calculated using student T-test. **p*< 0.05, ***p*< 0.05, ****p*< 0.001.

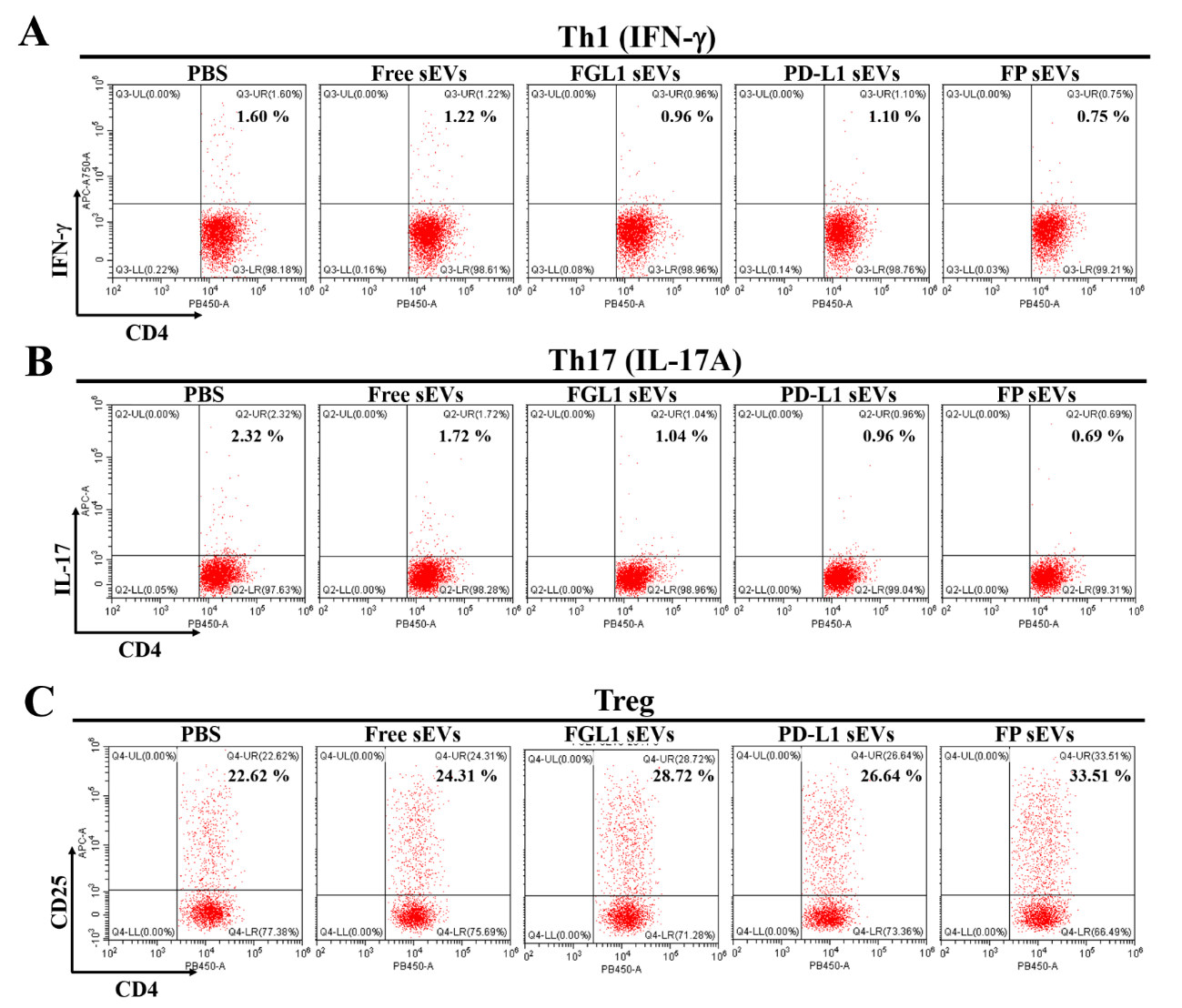
**Figure S6. FGL1/PD-L1 sEVs could affect Th1, Th17 and Treg differentiation.** (related to Figure 4)

A-C) Flow cytometry analysis of CD4^+^ IFN-γ^+^ Th1 (A), CD4^+^ IL-17A^+^ Th17 (B) and CD4^+^ CD25^+^ Treg (C) expression in P/I-stimulated CD3^+^ T cells treated with or without different groups of sEVs.

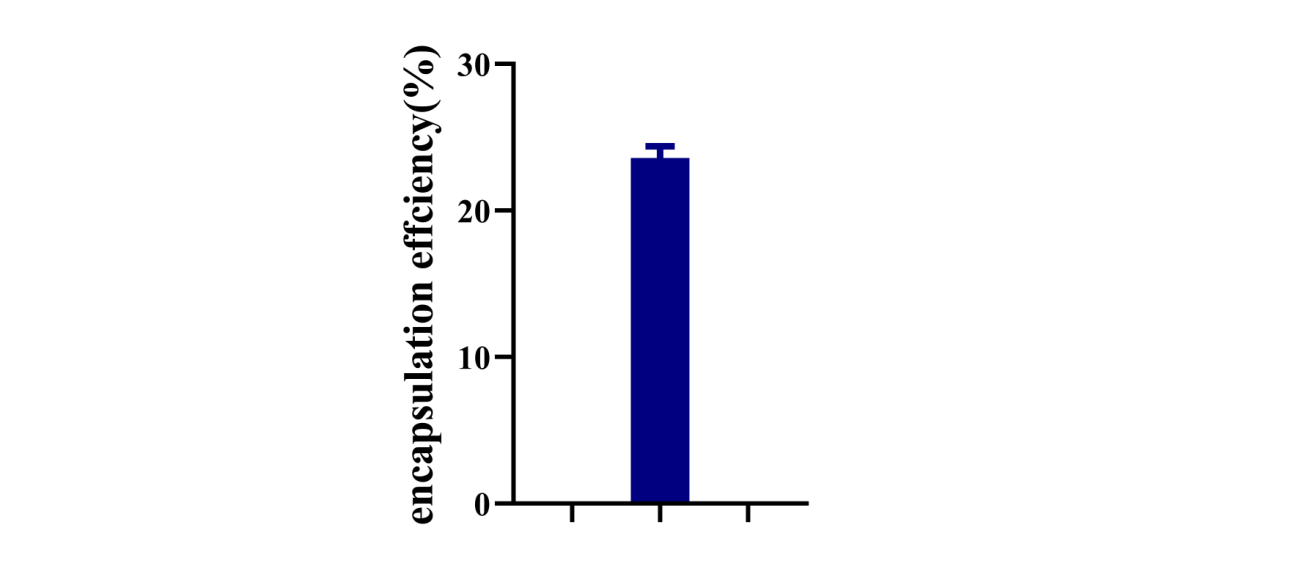

**Figure S7. Encapsulation rate of FGL1/PD-L1 sEVs loaded with FK506.** (related to Figure 4)

FK506 was encapsulated in the FGL1/PD-L1 sEVs by electroporation, and its encapsulation rate was detected by UV-VIS at 300 nm (*n* = 3).

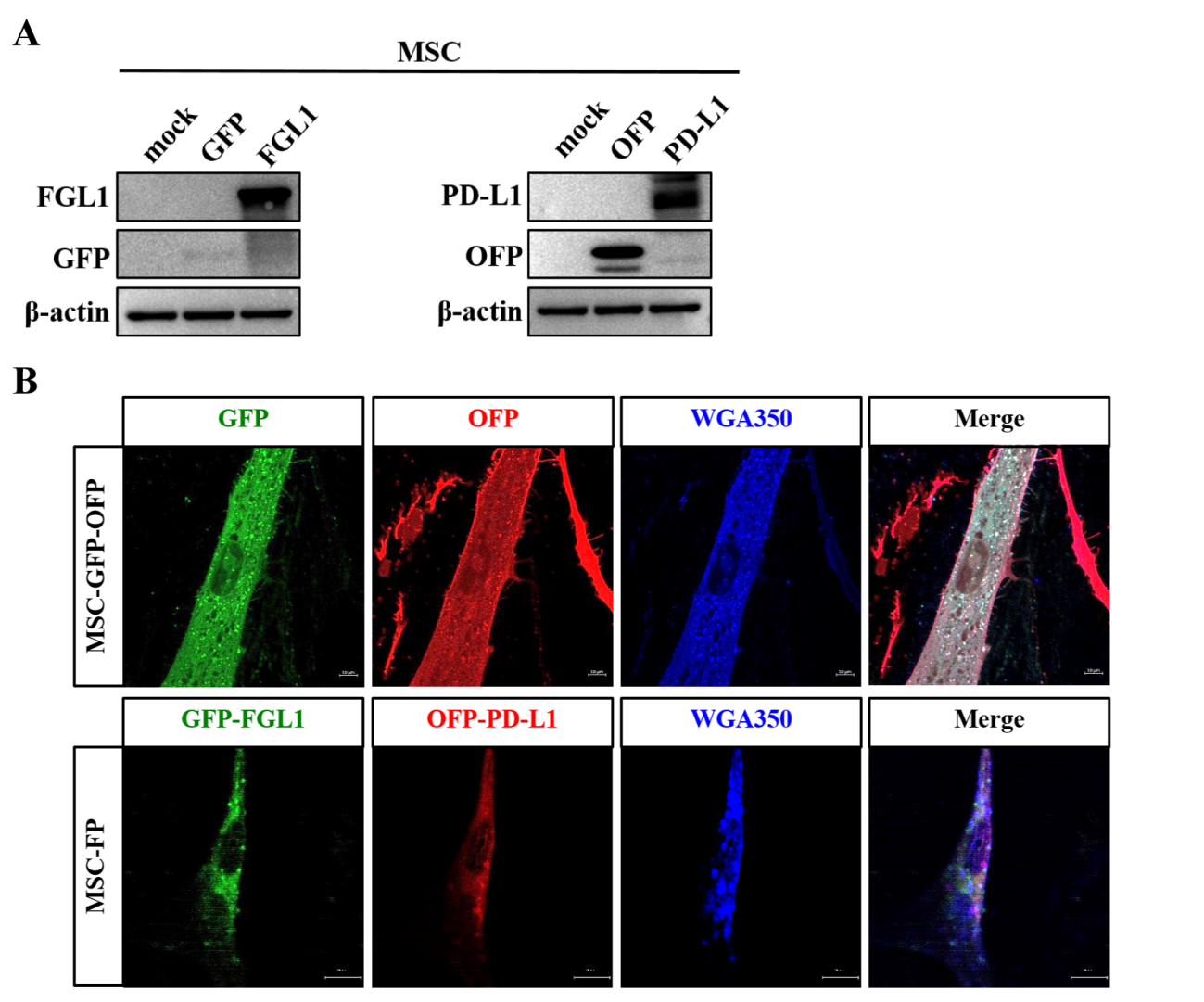

**Figure S8. Establishment of MSC cells expressing mouse-FGL1/PD-L1.** (related to Figure 5)

A) Western blotting for mouse-PD-L1-OFP and mouse-FGL1-GFP in the whole cell lysate in MSCs. B) Confocal images indicate the expression of mouse-PD-L1-OFP, mouse-FGL1-GFP and mouse-PD-L1-OFP/mouse-FGL1-GFP in MSCs. Cell membranes stained with WGA Alexa 350. Scale bar: 10 *µ*m.

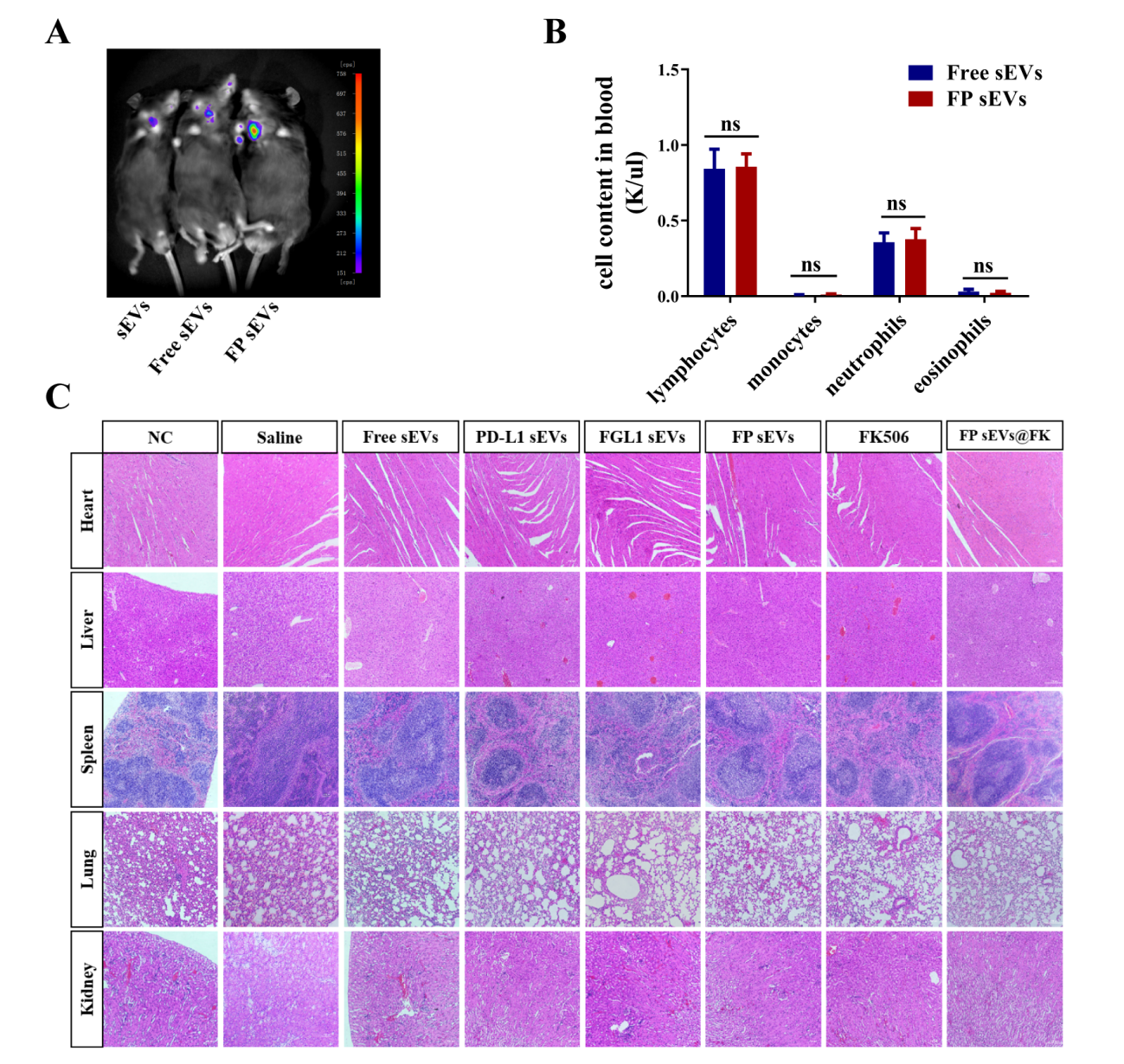

**Figure S9. Bioluminescence imaging and toxicity test of FGL1/PD-L1 sEVs in mice.** (related to Figure 5)

A) *In vivo* Bioluminescence imaging of sEVs, Free sEVs, and FP sEVs originating from MSC cells via tail vein injection. sEVs: injected with MSC-sEVs. Free sEVs: injected with vector sEVs. B) Complete blood count test (CBC test). Mice were injected with Free sEVs or FP sEVs via tail vein injection for 14 days for whole blood analysis, *n* = 5. Error bar, mean ± SEM. *P*-values are calculated using student T-test. ns: not significant. C) Histological images obtained from the hearts, livers, spleens and kidneys of mice treated with NC, saline, or different groups of EVs for 14 days post-injection as compared to the NC group. Scale bar: 100 *µ*m.

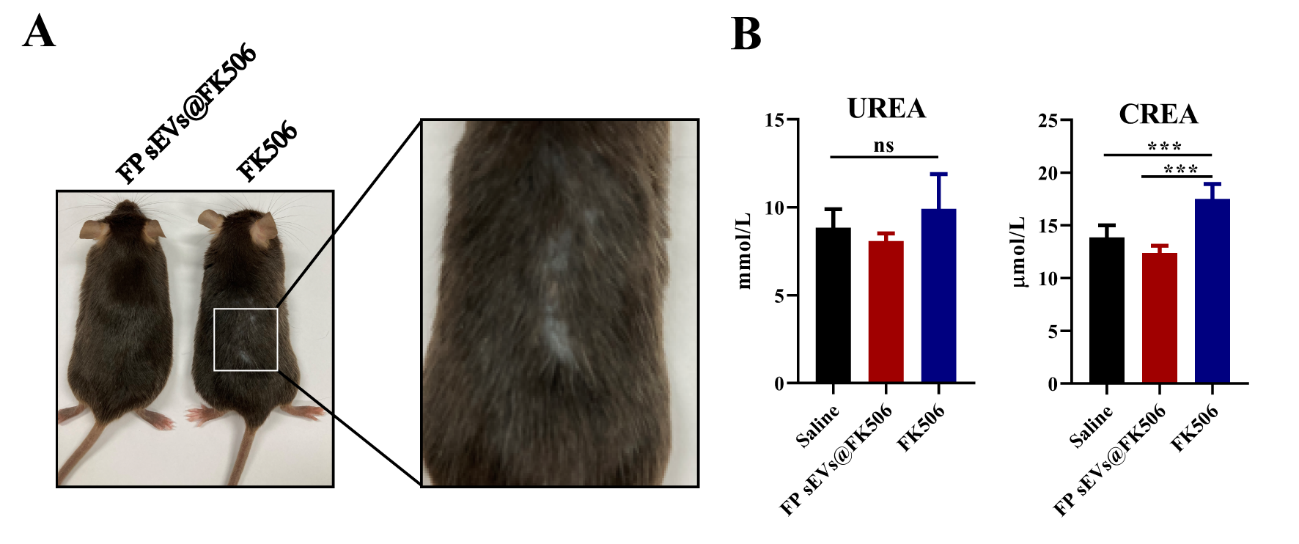

**Figure S10. The appearance and biochemical analysis of FK506 and FP sEVs @ FK506 in heart allogenic mice.** (related to Figure 5)

A) Images highlighting the hair loss of heart allogenic mice after injecting with FK506 or FP sEVs@FK506. B) Blood samples were collected after injecting with saline, FK506 or FP sEVs@FK506 to measure creatinine (CREA) and urea nitrogen (UREA) by automatic blood analyzer. *n* = 5. Error bar, mean ± SEM. *P < 0.05, **P < 0.01, ***P < 0.001.

**Supplementary Tables**

**Table S1. Top-scored negative immunoregulation miRNAs between HEK-293T sEVs and MSC-FP sEVs.**

| **Up-regulated miRNAs** | **ID** | ***P* VALUE** | **Function** |
| --- | --- | --- | --- |
| hsa-miR-199a-3p^[1]^ | MIMAT0000232 | 1.45713E-08 | Inhibition of inflammation, immunity and resistance to kidney damage |
| hsa-miR-23b-3p^[2]^ | MIMAT0000418 | 2.43588E-05 | Effective treatment of rheumatoid arthritis |
| hsa-miR-125a-5p^[3]^ | MIMAT0000443 | 0.008456264 | Promote the generation of M2 type macrophages |
| hsa-miR-3168^[4]^ | MIMAT0015043 | 8.46957E-25 | Suppress the immune system and relieve asthma |
| hsa-let-7i-5p^[5]^ | MIMAT0000415 | 1.19388E-33 | Inhibition of IL - 6 |
| hsa-miR-21-5p^[6]^ | MIMAT0000076 | 0.000657411 | Inhibits the immune function of dendritic cells |
| hsa-miR-23a-3p^[7]^ | MIMAT0000078 | 3.35175E-05 | Upregulate the expression of PD-L1 to inhibit T cells function |
| hsa-let-7b-5p^[8]^ | MIMAT0000063 | 4.70362E-27 | Inhibit the expression of inflammatory cytokines in macrophages |
| hsa-miR-125b-5p^[9]^ | MIMAT0000423 | 1.37316E-88 | Inhibition of inflammation, immunity and resistance to kidney damage |
| **Down-regulated miRNAs** | **ID** | ***P* VALUE** | **Function** |
| hsa-miR-4442^[10]^ | MIMAT0018960 | 1.2899E-122 | Downregulation inhibits inflammation and relieves dermatomyositis |
| hsa-miR-1273h-5p^[11]^ | MIMAT0030415 | 1.0546E-166 | Increased in the PBMCs of SLE patients |

**Table S2. Description of T cell related factors**

| **Factor** | **Function description** | **Reference** |
| --- | --- | --- |
| IL-2 | Promote the differentiation of CD4^+^T cells and activate Treg cells | ^[12]^ |
| IL-18 | Promote the differentiation of CD4^+^T cells | ^[13]^ |
| IL-22 | Promote the differentiation of CD4^+^T cells | ^[14]^ |
| IL-10 | Activate Treg cells | ^[15]^ |
| IL-5 | Regulating the differentiation, growth and activation of eosinophils | ^[16]^ |
| IL-17A | Promote the differentiation of CD4^+^T and CD8^+^ cells | ^[14, 17]^ |
| IL-21 | Inhibit the proliferation of Treg cells | ^[18]^ |
| IL-12 | Promote the differentiation of CD4^+^T cells | ^[18a, 19]^ |
| IL-6 | Regulator of Treg/Th17 balance | ^[20]^ |
| CCL1 | Suppressive function of Tregs | ^[21]^ |
| ID3 | Control of the differentiation of regulatory T cells and TH17 cells | ^[22]^ |
| ID1 | Promotes T Regulatory Cell Differentiation | ^[23]^ |
| TNFRSF12 | Promotes T Regulatory Cell Differentiation | ^[24]^ |
| BMPR1B | Development and Activation of CD4^+^ T Cells | ^[25]^ |
| CX3CL1 | Chemotactic activity for monocytes, NK cells, and T cells | ^[26]^ |
| IL11 | Promote the proliferation and differentiation of CD4^+^T cells | ^[27]^ |
| ILF | Regulator of Treg/Th17 balance | ^[28]^ |
| CXCL13 | Activate Tregs cells | ^[29]^ |
| CCL4 | Inhibit the proliferation of CD8^+^ cells | ^[30]^ |
| IL12RB2 | Promotes Th1 Cell Differentiation | ^[31]^ |
| TNFSF14 | Activate NK T cells | ^[32]^ |
| IDO1 | Activate Tregs cells | ^[33]^ |
| IL36A | Activate T cells | ^[34]^ |
| IL1RN | Activate Th17 cells | ^[35]^ |
| IL12A | Promotes CD8^+^ Cell Differentiation | ^[36]^ |
| IL21 | Promotes CD8^+^ Cell Differentiation | ^[37]^ |
| TNFRSF8 | Promotes Th17 Cell Differentiation | ^[38]^ |
| TNFRSF12A | Promotes the proliferation of T cells | ^[39]^ |
| EDA2R | Inhibit the Differentiation of Th17 cells | ^[40]^ |
| TSLP | enhances the function of helper type 2 cells | ^[41]^ |

**Table S3. The demographic clinical characteristics in kidney transplant patients.**

| **Variables** | **Total (n = 19)** | **ABMR (n = 6)** | **TCMR (n = 6)** | **Stable (n = 7)** | ***P*** |
| --- | --- | --- | --- | --- | --- |
| Donors |  |  |  |  |  |
| Types, n (%) |  |  |  |  | **0.045** |
| DD | 11 (58) | 2 (33) | 6 (100) | 3 (43) |  |
| LD | 8 (42) | 4 (67) | 0 (0) | 4 (57) |  |
| Recipients |  |  |  |  |  |
| Gender, n (%) |  |  |  |  | **0.037** |
| Female | 9 (47) | 1 (17) | 2 (33) | 6 (86) |  |
| Male | 10 (53) | 5 (83) | 4 (67) | 1 (14) |  |
| Age (year), Mean ± SEM | 36.96 ± 9.30 | 42.10 ± 10.44 | 37.43 ± 10.60 | 32.14 ± 4.61 | 0.156 |
| Weight (kg), Mean ± SEM | 52.0 ± 12.2 | 59.3 ± 16.1 | 50.2 ± 7.2 | 47.3 ± 10.2 | 0.191 |
| Previous_transplantation, n (%) |  |  |  |  | 0.48 |
| 0 | 16 (84) | 6 (100) | 4 (67) | 6 (86) |  |
| 1 | 3 (16) | 0 (0) | 2 (33) | 1 (14) |  |
| Days_from_transplant_to_sample, Median (IQR) | 371 (185, 582) | 1474 (822, 2142) | 133 (73, 152) | 376 (369, 378) | **0.002** |
| CNI^*^, n (%) |  |  |  |  | 1 |
| FK506 | 19 (100) | 6 (100) | 6 (100) | 7 (100) |  |
| FK506 C_0_ (ng/ml), Median (IQR) | 6.5 (5.10, 7.80) | 7.8 (7.58, 9.75) | 5.5 (5.05, 6.30) | 6.3 (5.55, 8.25) | 0.168 |
| White blood cell count (10^9/L), Median (IQR) | 7.44 ± 2.14 | 7.63 ± 2.82 | 7.71 ± 2.25 | 7.05 ± 1.63 | 0.847 |
| Lymphocyte count (10^9/L), Mean ± SEM | 1.35 ± 0.63 | 1.07 ± 0.38 | 1.32 ± 0.66 | 1.61 ± 0.75 | 0.331 |
| Neutrophil (10^9/L), Median (IQR) | 5.41 ± 2.12 | 5.83 ± 2.63 | 5.70 ± 1.71 | 4.80 ± 2.14 | 0.657 |
| Monocyte count (10^9/L), Mean ± SEM | 0.57 ± 0.17 | 0.62 ± 0.17 | 0.58 ± 0.22 | 0.52 ± 0.14 | 0.608 |

CNIs, Calcineurin inhibitors; FK506, Tacrolimus; LD, Living donor; DD, Death donor. ^*^ Standard maintenance immunosuppression, including CNI, prednisolone and mycophenolic acid was applied to all patients.

**Table S4. qPCR primer sequences.**

| **Gene** | **Forward primer sequence 5’→3’** | **Reverse primer sequence 5’→3’** |
| --- | --- | --- |
| mouse-*β-Actin* | GGCTGTATTCCCCTCCATCG | CCAGTTGGTAACAATGCCATGT |
| mouse-*Tnfα* | TCTCATCAGTTCTATGGCCC | GGGAGTAGACAAGGTACAAC |
| mouse-*Granzyme B* | TCGACCCTACATGGCCTTAC | TGGGGAATGCATTTTACCAT |
| mouse-*Pd-l1* | TCTGATCGTCGATTGGCAGC | CGTTGTTCCAGGCTCCTCTC |
| mouse-*Fgl1* | CTTCGTCCTGGTCGCCATT | TCCCGCAAGCAGTTCTCAC |
| mouse-*Tim3* | TCAGGTCTTACCCTCAACTGTG | GGCATTCTTACCAACCTCAAACA |
| mouse-*Vista* | GACAGGTGGCCTCTCACC | TTTTCGATTCCTTGGGTGTT |
| mouse-*Ctla4* | GCTTCCTAGATTACCCCTTCTGC | CGGGCATGGTTCTGGATCA |
| mouse-*Tigit* | CTGATACAGGCTGCCTTCCT | TGGGTCACTTCAGCTGTGTC |
| mouse-*Il-12* | CAGCATGTGTCAATCACGCTAC | TGTGGTCTTCAGCAGGTTT |
| mouse-*Pd-1* | ACCCTGGTCATTCACTTGGG | CATTTGCTCCCTCTGACACTG |
| mouse-*LAG-3* | AGGTTCCAGGGTGGGGC | GTCCACTTGGCAGTGAGGAA |
| mouse-*Foxp3* | CACCTATGCCACCCTTATCCG | CATGCGAGTAAACCAATGGTAGA |
| human-*LAG-3* | TCACTGTTCTGGGTCTGGAG | GGTAAAGTCGCCATTGTCTC |
| human-*PD-1* | GCACGAGGGACAATAGGAGCC | AATGGTGGCATACTCCGTCTG |
| human-*ACTIN* | CCACACTGTGCCCATCTAC | AGGATCTTCATGAGGTAGTCAGTC |
| human-*FGL1* | AGTCTGCTTCTGCTACTTGG | TCCCTCCATCTTCACCCTTATT |
| human-*PD-L1* | GGTGAGGATGGTTCTACACAG | GAGAACTGCATGAGGTTGC |
| human-*TBET* | GCAGCACCGCTACTTCTACC | GTAGGCGTAGGCTCCAAGG |
| human-*GATA3* | AGAGCGTGGCCTGGAGAC | CTCCCACAGTCAAGGAAACC |
| human-*FOXP3* | GAGAAGCTGAGTGCCATGCA | TTGATCTTGAGGTCAGGGGCCAGG |
| human-*RORγt* | AAATCTGTGGGGACAAGTCG | CTGACGGGTGCAGGAGTAG |
| human-U6 | CTCGCTTCGGCAGCACA | AACGCTTCACGAATTTGCGT |
| mQ-PRIMER-R | GTGCAGGGTCCGAGGT |  |
| miR-let-7b-5p-F | GCGCTGAGGTAGTAGGTTGT |  |
| miR-21-5p-F | GCGCTAGCTTATCAGACTGA |  |
| miR-125b-5p-F | GCTCCCTGAGACCCTAAC |  |

**Reference**

[1]	G. Zhu, L. Pei, F. Lin, H. Yin, X. Li, W. He, N. Liu, X. Gou, *J. Cell Physiol.* **2019**, *234* (12), 23736.

[2]	R. Li, Q. Ruan, F. Yin, K. Zhao, *J. Pharmacol. Sci.* **2021**, *145* (1), 69.

[3]	S. Banerjee, H. Cui, N. Xie, Z. Tan, S. Yang, M. Icyuz, V. J. Thannickal, E. Abraham, G. Liu, *J. Biol. Chem.* **2013**, *288* (49), 35428.

[4]	T. Bahmer, S. Krauss-Etschmann, D. Buschmann, J. Behrends, H. Watz, A. M. Kirsten, F. Pedersen, B. Waschki, O. Fuchs, M. W. Pfaffl, E. von Mutius, K. F. Rabe, G. Hansen, M. V. Kopp, I. R. Konig, S. Bartel, *Allergy* **2021**, *76* (1), 366.

[5]	X. Wang, H. X. Wang, Y. L. Li, C. C. Zhang, C. Y. Zhou, L. Wang, Y. L. Xia, J. Du, H. H. Li, *Hypertension* **2015**, *66* (4), 776.

[6]	M. Reis, E. Mavin, L. Nicholson, K. Green, A. M. Dickinson, X. N. Wang, *Front. Immunol.* **2018**, *9*, 2538.

[7]	J. Liu, L. Fan, H. Yu, J. Zhang, Y. He, D. Feng, F. Wang, X. Li, Q. Liu, Y. Li, Z. Guo, B. Gao, W. Wei, H. Wang, G. Sun, *Hepatology* **2019**, *70* (1), 241.

[8]	S. E. Nematian, R. Mamillapalli, T. S. Kadakia, M. Majidi Zolbin, S. Moustafa, H. S. Taylor, *J. Clin. Endocrinol. Metab.* **2018**, *103* (1), 64.

[9]	J. Y. Cao, B. Wang, T. T. Tang, Y. Wen, Z. L. Li, S. T. Feng, M. Wu, D. Liu, D. Yin, K. L. Ma, R. N. Tang, Q. L. Wu, H. Y. Lan, L. L. Lv, B. C. Liu, *Theranostics* **2021**, *11* (11), 5248.

[10]	Y. Y. Yang, X. X. Zuo, H. L. Zhu, S. J. Liu, *Beijing Da Xue Xue Bao Yi Xue Ban* **2019**, *51* (2), 374.

[11]	G. Guo, H. Wang, X. Shi, L. Ye, K. Wu, K. Lin, S. Ye, B. Li, H. Zhang, Q. Lin, S. Ye, X. Xue, C. Chen, *J. Transl. Med.* **2018**, *16* (1), 370.

[12]	a) H. Abken, *Transplantation* **2021**, *105* (7), 1394; b) S. Sad, T. R. Mosmann, *J. Immunol.* **1994**, *153* (8), 3514.

[13]	M. Sawada, T. Kawayama, H. Imaoka, Y. Sakazaki, H. Oda, S. Takenaka, Y. Kaku, K. Azuma, M. Tajiri, N. Edakuni, M. Okamoto, S. Kato, T. Hoshino, *PLoS One* **2013**, *8* (1), e54623.

[14]	T. J. Scriba, B. Kalsdorf, D. A. Abrahams, F. Isaacs, J. Hofmeister, G. Black, H. Y. Hassan, R. J. Wilkinson, G. Walzl, S. J. Gelderbloem, H. Mahomed, G. D. Hussey, W. A. Hanekom, *J. Immunol.* **2008**, *180* (3), 1962.

[15]	a) J. Niu, W. Yue, Y. Song, Y. Zhang, X. Qi, Z. Wang, B. Liu, H. Shen, X. Hu, *Clin. Exp. Immunol.* **2014**, *176* (3), 473; b) C. I. Kingsley, M. Karim, A. R. Bushell, K. J. Wood, *J. Immunol.* **2002**, *168* (3), 1080.

[16]	M. Goldman, A. Le Moine, M. Braun, V. Flamand, D. Abramowicz, *Trends in immunology* **2001**, *22* (5), 247.

[17]	M. P. Crawford, S. Sinha, P. S. Renavikar, N. Borcherding, N. J. Karandikar, *Proc. Natl. Acad. Sci. U. S. A.* **2020**, *117* (32), 19408.

[18]	a) C. Jandl, S. M. Liu, P. F. Canete, J. Warren, W. E. Hughes, A. Vogelzang, K. Webster, M. E. Craig, G. Uzel, A. Dent, *Nat. Commun.* **2017**, *8* (1), 1; b) A. Suto, D. Kashiwakuma, S. Kagami, K. Hirose, N. Watanabe, K. Yokote, Y. Saito, T. Nakayama, M. J. Grusby, I. Iwamoto, H. Nakajima, *J. Exp. Med.* **2008**, *205* (6), 1369.

[19]	G. Trinchieri, S. Pflanz, R. A. Kastelein, *Immunity* **2003**, *19* (5), 641.

[20]	A. Kimura, T. Kishimoto, *Eur. J. Immunol.* **2010**, *40* (7), 1830.

[21]	D. B. Hoelzinger, S. E. Smith, N. Mirza, A. L. Dominguez, S. Z. Manrique, J. Lustgarten, *J. Immunol.* **2010**, *184* (12), 6833.

[22]	T. Maruyama, J. Li, J. P. Vaque, J. E. Konkel, W. Wang, B. Zhang, P. Zhang, B. F. Zamarron, D. Yu, Y. Wu, Y. Zhuang, J. S. Gutkind, W. Chen, *Nat. Immunol.* **2011**, *12* (1), 86.

[23]	C. Liu, H. C. Wang, S. Yu, R. Jin, H. Tang, Y. F. Liu, Q. Ge, X. H. Sun, Y. Zhang, *J. Immunol.* **2014**, *193* (2), 663.

[24]	H. Nishikii, B. S. Kim, Y. Yokoyama, Y. Chen, J. Baker, A. Pierini, M. Alvarez, M. Mavers, K. Maas-Bauer, Y. Pan, S. Chiba, R. S. Negrin, *Blood* **2016**, *128* (24), 2846.

[25]	M. Kuczma, P. Kraj, *Vitam. Horm.* **2015**, *99*, 171.

[26]	B. A. Jones, M. Beamer, S. Ahmed, *Mol. Interv.* **2010**, *10* (5), 263.

[27]	A. Curti, A. Tafuri, M. R. Ricciardi, P. Tazzari, M. T. Petrucci, M. Fogli, M. Ratta, R. Lapalombella, E. Ferri, S. Tura, M. Baccarani, R. M. Lemoli, *Haematologica* **2002**, *87* (4), 373.

[28]	S. M. Metcalfe, *Genes Immun.* **2011**, *12* (3), 157.

[29]	B. P. Lee, W. Chen, H. Shi, S. D. Der, R. Forster, L. Zhang, *J. Immunol.* **2006**, *176* (9), 5276.

[30]	S. A. Joosten, K. E. van Meijgaarden, N. D. Savage, T. de Boer, F. Triebel, A. van der Wal, E. de Heer, M. R. Klein, A. Geluk, T. H. Ottenhoff, *Proc. Natl. Acad. Sci. U. S. A.* **2007**, *104* (19), 8029.

[31]	Z. Y. Zhou, S. L. Chen, N. Shen, Y. Lu, *Autoimmun Rev.* **2012**, *11* (10), 699.

[32]	F. Shi, Y. Xiong, Y. Zhang, C. Qiu, M. Li, A. Shan, Y. Yang, B. Li, *Inflammation* **2018**, *41* (3), 1021.

[33]	A. de Luca, S. Bozza, T. Zelante, S. Zagarella, C. D'Angelo, K. Perruccio, C. Vacca, A. Carvalho, C. Cunha, F. Aversa, L. Romani, *Cell Mol. Immunol.* **2010**, *7* (6), 459.

[34]	S. Abdollahi-Roodsaz, L. A. Joosten, M. I. Koenders, I. Devesa, M. F. Roelofs, T. R. Radstake, M. Heuvelmans-Jacobs, S. Akira, M. J. Nicklin, F. Ribeiro-Dias, W. B. van den Berg, *J. Clin. Invest.* **2008**, *118* (1), 205.

[35]	J. C. Waite, D. Skokos, *Int. J. Inflam.* **2012**, *2012*, 819467.

[36]	N. Takemoto, A. M. Intlekofer, J. T. Northrup, E. J. Wherry, S. L. Reiner, *J. Immunol.* **2006**, *177* (11), 7515.

[37]	C. S. Hinrichs, R. Spolski, C. M. Paulos, L. Gattinoni, K. W. Kerstann, D. C. Palmer, C. A. Klebanoff, S. A. Rosenberg, W. J. Leonard, N. P. Restifo, *Blood* **2008**, *111* (11), 5326.

[38]	X. Sun, H. Yamada, K. Shibata, H. Muta, K. Tani, E. R. Podack, Y. Yoshikai, *J. Immunol.* **2010**, *185* (4), 2222.

[39]	L. Xia, L. Jiang, Y. Chen, G. Zhang, L. Chen, *Cytokine* **2021**, 155658.

[40]	P. G. Miller, M. B. Bonn, S. C. McKarns, *J. Immunol.* **2015**, *195* (6), 2633.

[41]	M. Kitajima, H. C. Lee, T. Nakayama, S. F. Ziegler, *Eur. J. Immunol.* **2011**, *41* (7), 1862.
